# GeneSys: Generative Modeling of Developmental System

**DOI:** 10.1101/2025.08.20.671385

**Authors:** Che-Wei Hsu, Chia-Yu Chen, Trevor M. Nolan, Philip N. Benfey, Uwe Ohler

## Abstract

Temporal single-cell transcriptomics enables the reconstruction of dynamic gene expression changes during development. Yet, its analytical power is often limited by data sparsity, technical noise, and imbalanced representation of cell types across time points. To overcome these challenges, we present GeneSys (Generative Modeling of Developmental System), a generative deep learning model that simulates single-cell transcriptomic landscapes under developmental constraints, which is informed by prior biological knowledge or user-defined hypotheses. GeneSys integrates a temporal variational autoencoder with a cell-type classifier, requiring a lineage blueprint as input, which enables it to model the temporal transitions of transcriptional states with cell-type specificity. Leveraging data from *Arabidopsis thaliana* roots and mouse embryos, we show that GeneSys learns robust developmental trajectories, generates imputed and representative transcriptomes, and enhances gene prioritization accuracy compared to unregularized scRNA-seq data. By applying gene masking and augmentation, GeneSys reveals interpretable gene expression programs (GEPs) and serves as an *in silico* platform to test the impact of specific genes or gene sets on user-defined developmental outcomes. Additionally, GeneSys computes linear interaction matrices (LIMAs) to infer dynamic gene networks and prioritize transcription factors with spatiotemporal resolution. These features enable GeneSys to nominate key genes governing state transitions in a developmental system, supporting both mechanistic insight and hypothesis generation. Together, GeneSys provides a flexible and extensible framework to denoise single-cell data and simulate transcriptomic developmental landscape guided by known or hypothesized developmental constraints, empowering the discovery of regulatory mechanisms from high-dimensional single-cell datasets.

## Introduction

Single-cell RNA sequencing (scRNA-seq) has emerged as a powerful tool for capturing the transcriptomic profiles of individual cells, enabling researchers to identify and characterize diverse tissues and cell types^1^. By collecting scRNA-seq data across multiple time points or inferring the developmental trajectories of each cell type, researchers can trace the dynamics of transcriptomic changes at single-cell resolution over time^2, 3^.

Temporal information is crucial for studying various cellular processes, such as cell differentiation, cell cycle progression, responses to stimuli, host-microbes interaction, growth and development^4^. Sampling cells at different time points allows researchers to detect transient or rare cell states that might otherwise be overlooked in static, single-time-point analyses. Moreover, this temporal data aids in the construction of gene regulatory networks (GRNs), which can infer regulatory relationships by leveraging changes in gene expression at the single-cell level to reveal the underlying molecular mechanisms driving cellular, organ, and organismal behavior^5, 6^.

Despite the promising applications of temporal inference in single-cell transcriptomics, several challenges remain in the analysis and modeling of developmental dynamics. The inherent sparsity of scRNA-seq data often necessitates denoising and imputation to reduce uncertainty in downstream analyses^7, 8^. Combined with technical noise, bias and variation introduced during sample and library preparation, this can lead to the underrepresentation or overrepresentation of specific cell types and developmental stages^9, 10^. Such imbalances complicate the task of linking cells within and across time points, making it challenging to learn continuous trajectories for each cell type. As a result, researchers may struggle to obtain representative transcriptomic profiles across developmental time, leading to inferred GRNs that fail to fully capture the dynamics of developmental processes^9, 11^. Attempts to address this limitation by integrating scRNA-seq data from different studies to increase cell numbers for each time point introduce new challenges: batch effects and technical noise from diverse profiling platforms and different choice of preprocessing pipelines further hinder accurate reconstruction of cellular trajectories^9^.

To address these challenges, single-cell data scientists have used both statistical models and deep neural networks to denoise, impute, and integrate data. Either by modeling the distribution of noise and true biological signal or by training unsupervised autoencoders to learn robust data representations^9^. However, these approaches face persistent issues: model selection is often arbitrary and based on assumptions that may deviate from the ground truth, and reconstruction by neural networks could further exacerbate sampling biases toward abundant cell types or states introduced during sample and library preparation. We therefore argue that denoising and imputation should be guided by domain knowledge, developmental constraints, or specific hypotheses to minimize bias and ensure balanced representation of all cell states in a system. This highlights the need for computational techniques capable of encoding such knowledge directly into the imputation process. In addition, a typical developmental system encompasses both cell differentiation and cellular maturation, the two tightly coordinated processes that drive development. Consequently, imputation methods should explicitly model each of these aspects.

To meet the need, we developed GeneSys, a deep generative model trained on annotated scRNA-seq data that incorporates both cell type identities and temporal information, such as developmental stages or treatment time points in time-series datasets. GeneSys combines a temporal variational autoencoder with a cell type classifier and requires a user-defined cell lineage blueprint to configure the known or hypothesized developmental pathways for each cell type within a biological system. It learns the transcriptomic dynamics along these trajectories and generates representative single-cell transcriptomes for any specified cell state. This generative capability enables the extraction of gene expression dynamics that span developmental transitions, while also facilitating noise reduction and imputation of missing data. In essence, GeneSys allows researchers to encode developmental constraints directly into the learning process of latent features, thereby reconstructing the transcriptomic landscape of the system in a biologically informed manner.

In this study, we demonstrate the utility of GeneSys using scRNA-seq datasets from *Arabidopsis thaliana* roots^12^ and a mouse embryo developmental time series^13^, where carefully curated annotations for both the cell types and temporal information are available. The *Arabidopsis* root serves as a simple and tractable developmental system due to its spatial organization, where cell types are arranged in concentric layers on the radial axis and developmental progression from stem cells to differentiated tissue is present along the longitudinal axis^12, 14^. This spatial-temporal alignment reflects developmental pathways for each cell type and enables pseudotime inference from a single snapshot of scRNA-seq data, minimizing batch effects and eliminating the need for true time-series sampling. The availability of extensive resources, including validated markers and bulk expression profiles, provides high-quality annotations that support both training and validation of GeneSys.

To further evaluate generalizability of GeneSys, we applied it to study a representative time-series dataset of mouse embryo development, which provides comprehensive annotations and a relatively well-established cell lineage blueprint. Unlike the root system, mouse embryogenesis involves many cell types that do not emerge until later stages, requiring extensive encoding of developmental pathways into the lineage blueprint based on domain knowledge or hypothesis. Together, these two systems illustrate the broad applicability of GeneSys across diverse biological contexts.

## Results

### The GeneSys model

GeneSys comprises two major neural network components: a temporal difference variational auto-encoder (TD-VAE)^15^ structured upon a bi-directional LSTM (Long Short-Term Memory) network,^16, 17^ and a multi-label classifier (cell type classifier)^18^ (Figure 1A; Figure S1). The LSTM network here is designed to process time-series data, with each LSTM cell (computational cell) representing a time step. The interconnected LSTM cells enable the learning of features associated with order dependency, making the LSTM network suitable for simulating a developmental system with continuous developmental trajectories.

**Figure 1.**
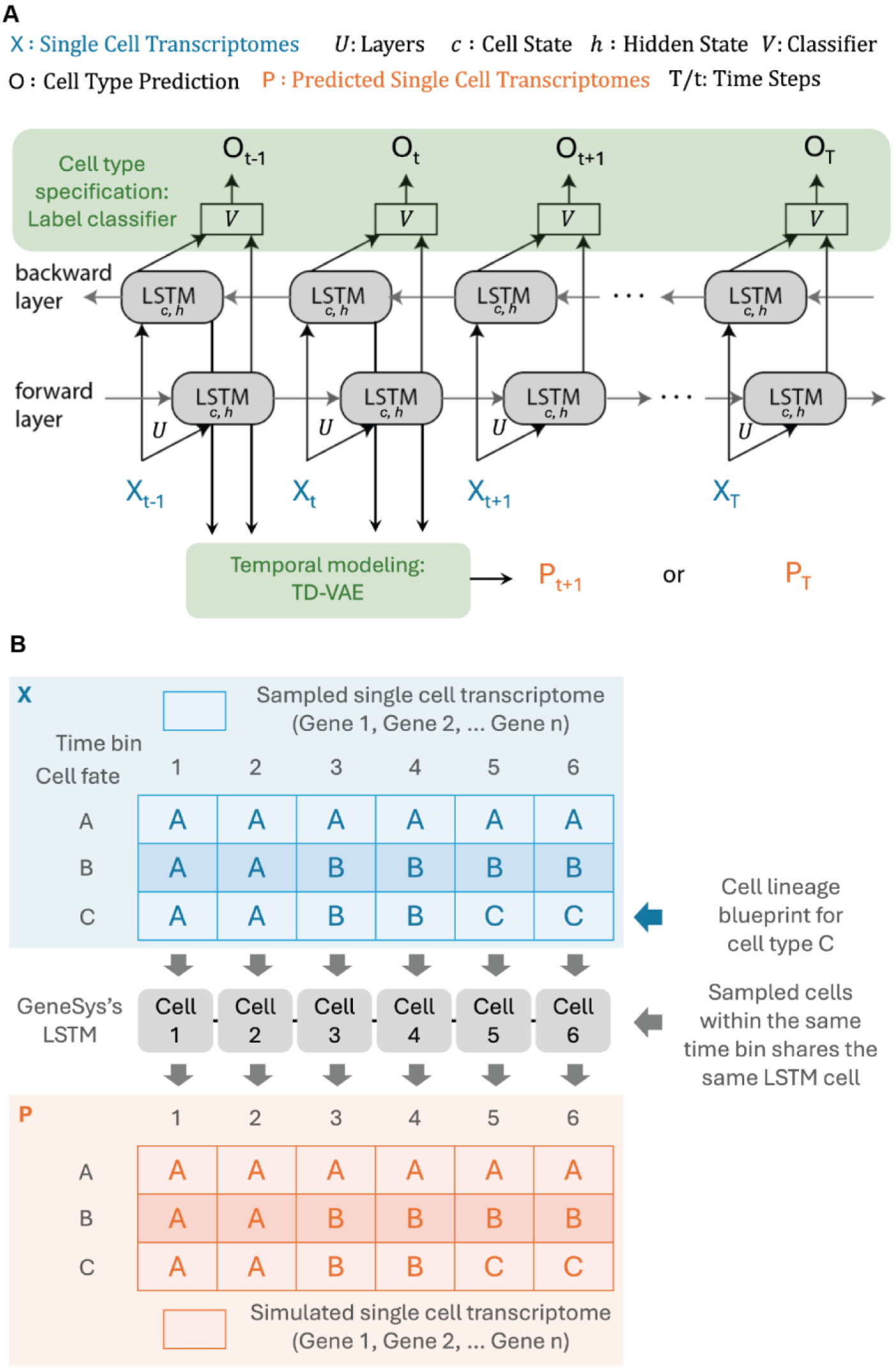
GeneSys model architecture. (A) Schematic of the GeneSys model. A bi-directional LSTM network is connected to both a multi-label cell type classifier and a temporal difference variational autoencoder (TD-VAE), enabling the model to learn cell type specificity and temporal developmental dynamics, respectively. Notation: **X** - single-cell transcriptomes; **U** - neural network layers; **c** - LSTM cell state; **h** – LSTM hidden state; **V** - cell type classifier; **O** - predicted cell type labels; **P** - predicted single-cell transcriptomes; **t** - specific LSTM time step; **T** - any LSTM time step. (B) Illustration of the relationship among training batches, the GeneSys LSTM, and the simulated transcriptomes. Each training batch is assembled based on a cell lineage blueprint, with single-cell transcriptomes randomly sampled from the corresponding cell types and time bins to form a training tensor. During training, transcriptomes from the same time bin share the same LSTM computational cell. The TD-VAE module of the trained model then generates a tensor of simulated transcriptomes that recapitulates the lineage blueprint.

In the GeneSys model, a user-defined cell lineage blueprint determines the architecture of the LSTM network, the output dimensions of the cell type classifier, and the sampling strategy used to construct training batches. Each LSTM cell corresponds to either a discrete time point (in time-series data) or a defined developmental stage, and the classifier’s output dimensions are determined by the number of distinct cell types in the training data.

Conceptually, GeneSys models developmental trajectories by leveraging the cell lineage blueprint, which describes how specific cell types emerge over time. For instance, in a system where cell type C arises from B, and B derives from A, the developmental trajectory of cell type C reflects this lineage: beginning with A, transitioning through B, and culminating in C. This lineage-based framework allows GeneSys to learn biologically grounded patterns of gene expression progression across developmental stages (Figure 1B). In cases where developmental lineages are not fully known, users can encode their best hypotheses or assumptions, with the understanding that the trained model will simulate trajectories under those specified conditions.

Training batches are assembled according to the specified cell lineage blueprint (see Materials and Methods), and are then fed into the LSTM network, where each time step corresponds to a developmental stage. As hidden states propagate through the TD-VAE, the model learns to generate transcriptomic profiles time steps ahead (including current time step) based on prior steps. Simultaneously, these hidden states are passed to the cell type classifier, enabling the model to predict the corresponding cell type for each trajectory. Training proceeds iteratively on such batches until the combined loss from the TD-VAE and the classifier converges (Figure S2). The resulting model generates transcriptomic profiles that, consistent with the properties of a variational autoencoder^8, 15^, are expected to be regularized, robust and representative of the cell type at each stage given the developmental relationships encoded in the lineage blueprint.

### GeneSys reconstructs root developmental trajectories *in silico*

We first applied GeneSys to scRNA-seq data from the *Arabidopsis* root atlas described by Shahan and Hsu et al. (2022)^12^. The transcriptomes from the root atlas were partitioned into three subsets: a training set (64% of cells) for model training, a validation set (16%) for tuning model parameters during training, and a test set (20%) for evaluating model performance. Cell type annotations and inferred pseudotime data were used to define developmental trajectories composed of eleven time bins. The root cell lineage blueprint includes ten major cell types and spans the eleven pseudotime bins, with quiescent center (QC) cells designated as the first time bin (t0) (Figure S3A).

Each cell of the root atlas was profiled for 17513 genes, allowing a single developmental trajectory to be represented as a matrix of shape (11, 17513), corresponding to eleven pseudotime bins. Given that the atlas includes ten major cell types, the training batches of shape (10, 11, 17513) are assembled, one trajectory per cell type with corresponding single cell transcriptomes randomly sampled from the training set according to the defined cell lineage blueprint (Figure S3A). During training, GeneSys’s LSTM network operates over a sequence length of 11, with each LSTM cell corresponding to one time bin (Figure 1B). The hidden states produced by the LSTM are passed to both a multi-label classifier (to learn cell type specificity) and a TD-VAE (to model temporal maturation dynamics).

After training, GeneSys was used to generate simulated transcriptomes from test batches spanning complete developmental trajectories (Figure 2A). The fidelity of these simulations was validated by the presence of known marker genes—such as GL2^19^ (atrichoblast), COBL9^20^ (trichoblast), CORTEX/AED3^21^ (cortex), MYB36^22^ (endodermis), APL^23^ (phloem), and VND7/NAC030^24^ (xylem) in the generated trajectories, which closely resembled those in the root atlas (Figure 2B). Compared to the original test batch inputs, the model-generated transcriptomes exhibited smoother and more continuous trajectories, with dense and regularized gene expression profiles (Figure 2A, 2C; Figure S4).

**Figure 2.**
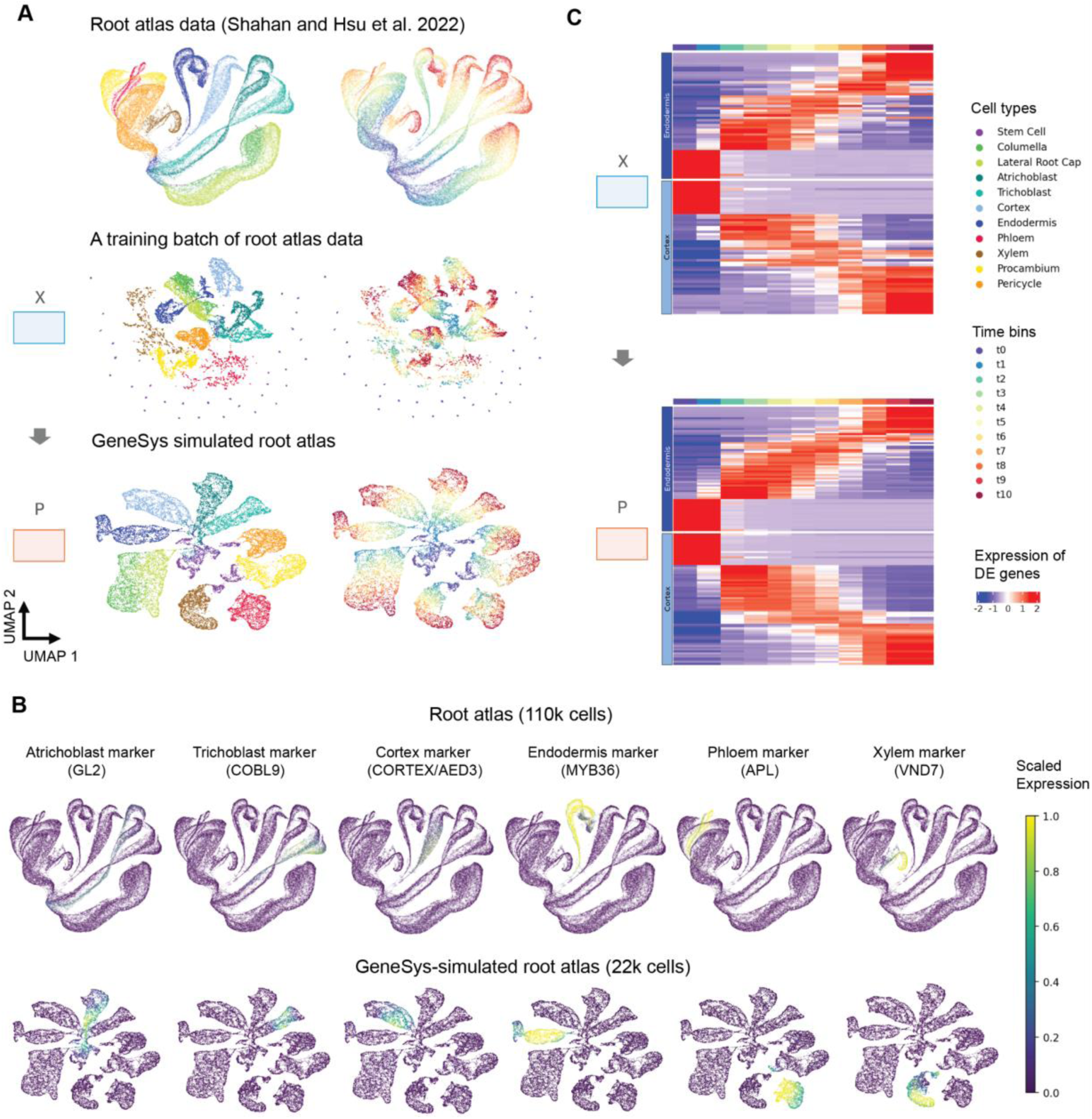
The trained GeneSys model accurately simulates the root atlas and generates dense, regularized single-cell transcriptomes. (A) UMAP embeddings of the original root atlas, a representative training batch X, and the GeneSys-simulated root atlas P. The training batch X and the simulated data P contain the same number of cells. (B) UMAPs showing the expression of known marker genes in the original and simulated root atlas, highlighting the preservation of cell type-specific expression patterns. (C) Scaled expression heatmaps of non-redundant differentially expressed (DE) genes across eleven pseudotime bins for cortex and endodermis. Comparisons are shown between the training input X and the simulated output P.

To probe the developmental representations learned by GeneSys, we assessed how many time bins could be masked from test inputs while still allowing the model to accurately reconstruct complete developmental trajectories. To quantify this, we introduced a metric termed recreation accuracy (see Materials and Methods), which measures the model’s ability to recover cell identities and generate representative transcriptomes for future time bins based on partially masked inputs. Using previously unseen wild-type single-cell data, we found that GeneSys achieved recreation accuracies of 0.75 and 0.88 when provided with only two or three input time bins, respectively (Figure S5A). The first two bins (t0 and t1) correspond to the proliferation domain of the root meristem, while the third (t2) represents the early transition domain^25^ (Figure S5B). These results suggest that GeneSys effectively models the processes of cell type specification and maturation during root development, but requires at least one to two early time point transitions as priming clues to accurately reconstruct full trajectories.

### GeneSys reveals fate-associated gene expression programs through gene set masking and augmentation

When supplied with all time bins, the trained GeneSys model functions as a virtual developmental system, enabling *in silico* testing of the impact of specific genes or gene sets on cell fate outcomes. For example, given the expression profiles of selected genes, one can estimate their contributions to cell fate using a metric that we devised, termed the *cell type recovery rate*, which reflects how accurately GeneSys can predict terminal cell identities using only those genes in masked test batches.

To evaluate the model’s dependency on the scale of gene expression input, we examined how the cell type recovery rate changed with decreasing numbers of genes. Remarkably, GeneSys achieved a perfect average recovery rate (1.0) even when using only 10% of randomly selected genes (1751 genes, averaged over 10 trials), comparable to the performance when all genes were used (Figure 3). This finding highlights the redundancy of gene expression information, likely due to co-expression of genes in functional modules. As randomly selected input gene counts dropped to 5%, 3%, and 1% (876, 525, and 175 genes, respectively), average recovery rates declined to 0.94, 0.71, and 0.17. These results indicate that just 5-10% of the transcriptome, randomly sampled, is sufficient to recapitulate the developmental landscape of the *Arabidopsis* root.

**Figure 3.**
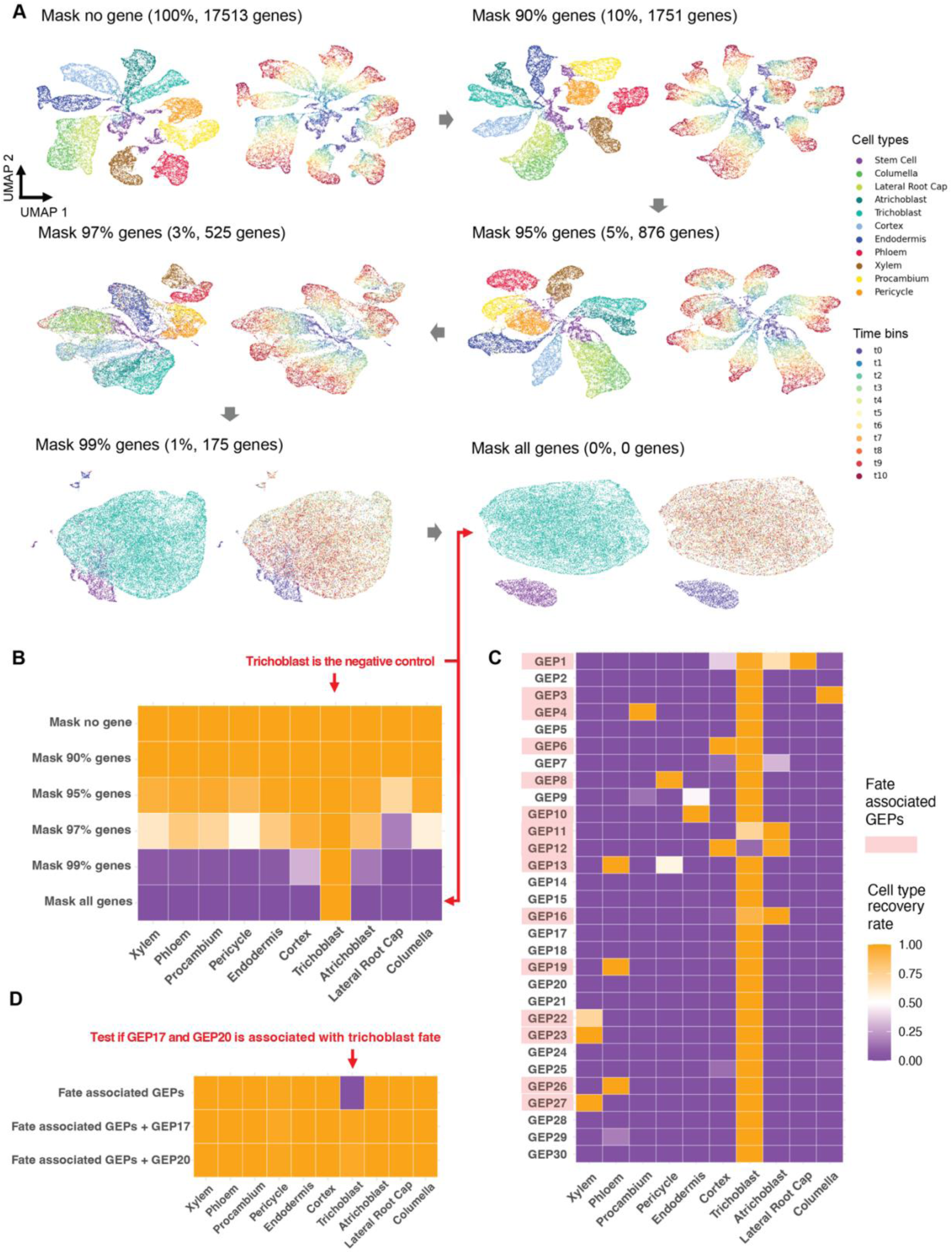
GeneSys-predicted developmental outcomes of gene masking and gene expression programs (GEPs) in the root. (A) UMAPs of GeneSys-simulated transcriptomes generated with varying percentages of randomly masked input genes. (B) Cell type recovery rates predicted by the trained GeneSys model under increasing levels of random gene masking. When all genes are masked, trichoblast consistently emerges as the default predicted cell type, serving as a negative control or baseline for assessing the developmental impact of specific gene sets. (C) Cell type recovery rates based on the top 30 representative genes from each of the 31 gene expression programs (GEPs) identified in the root atlas. GEPs shaded in red indicate those associated with cell fate, as determined by elevated recovery rates of non-trichoblast cell types. (D) Leave-one-out validation for GEPs suspected to be associated with trichoblast fate. Recovery rates are compared before and after reintroducing the candidate GEPs.

Based on this observation, we hypothesized that GeneSys may implicitly encode gene expression programs (GEPs), conceptually related to clusters of co-expressed genes or functional gene modules, that represent the developmental knowledge captured by the model.^26–28^ To explore this, we applied consensus non-negative matrix factorization (cNMF)^26^ to identify 30 GEPs in the root atlas, each comprising a set of genes exhibiting coordinated behavior across cells or conditions (Figure S6). We annotated each GEP by assessing its expression specificity across cell types and developmental stages, supplemented by gene ontology predictions. This analysis revealed two broad GEP categories: (1) cell type-specific programs likely involved in fate determination, and (2) broadly expressed programs likely reflecting core biological processes such as biotic and abiotic stress responses, cell cycle phases, and active translation (Table S1, Figure S7-9).

To quantify the functional relevance of each GEP, we calculated cell type recovery rates using only the top 30 representative genes (genes with the highest gene-by-factor loadings) from each GEP as model input. As a negative control, we masked all gene expression inputs, which resulted in an average recovery rate of 0.1, with all outputs predicted as trichoblasts (1.0 for trichoblast; 0 for all other cell types; Figure 3A, 3B). Against this baseline, several GEPs achieved recovery rates exceeding 0.7 for specific cell types, underscoring their critical roles in fate specification (Figure 3C). These results were well aligned with GEP annotations based on cell type specific expression and ontology.

Notably, both GEP17 and GEP20 were annotated as trichoblast-associated based on the gene ontology analysis and expression specificity. However, their roles in trichoblast fate specification could not be directly verified using cell type recovery rate alone, due to the negative control bias where all cells defaulted to trichoblast. To address this, we performed a leave-one-out experiment: we first included all fate-associated GEPs except GEP17 and GEP20, which resulted in cell fate recovery for all cell types except trichoblast. Reintroducing either GEP17 or GEP20 restored trichoblast recovery to high levels, suggesting they are important in trichoblast fate determination (Figure 3D).

In summary, the trained GeneSys model, combined with cell type recovery rate estimation, enhances both the resolution and annotation quality of the identified GEPs. It robustly characterizes 17 cell fate-associated GEPs with high cell type specificity, offering a valuable reference for future efforts in GEP engineering.

### TF prioritization efficiency is enhanced with GeneSys-generated profiles over scRNA-seq data

Since GEPs represent clusters of gene modules with shared biological functions, identifying the regulators that orchestrate each GEP is critical for understanding and potentially controlling developmental and physiological processes. Transcription factors (TFs) often serve this role by activating or repressing GEPs that govern cell fate decisions and tissue-specific functions. As such, identifying key TFs, or revealing novel roles for known ones, is a central goal in many single-cell studies.

To support this, a robust and accurate gene prioritization strategy is essential for guiding functional validation. We evaluated two main approaches: 1) Differential expression (DE) analysis, which ranks genes by fold-change, difference in expression proportions, or statistical significance across cell types or stages. 2) Network-based centrality analysis, which prioritizes TFs based on their influence within a given gene network. Centrality metrics, including in-degree, out-degree, betweenness centrality, and eigenvector centrality, provide complementary insights into regulatory importance by capturing network connectivity and information flow.

With a generative model such as GeneSys, gene to gene relationships can be inferred directly from transitions between cell-by-gene expression matrices. To capture this, we computed Linear Interaction Matrices (LIMAs) (see Materials and Methods), which model how changes in gene expression in one state linearly influence those in which it is about to become. These matrices can be converted into directed gene interaction networks, enabling TF prioritization based on interaction strength and directionality.

We evaluated three categories of TF prioritization strategies: (1) differential expression (DE) analysis, (2) centrality scores derived from LIMAs, and (3) centrality scores from GRNs inferred using CellOracle^5, 25^. Each strategy was applied to both the annotated root atlas scRNA-seq data and GeneSys-generated transcriptomes. Performance was assessed using a gold-standard set of 140 TFs, including tissue- and cell-type–specific subsets (see Materials and Methods).

Effectiveness was quantified using the R50 metric^29^, which measures the rank at which 50% of true positive TFs (TFs in the gold standard) appear in the prioritized list (a given gene ranking). Lower R50 values indicate better performance of the ranking scheme, as relevant TFs are ranked higher in the gold standard.

We tested twelve strategies: two based on DE, five using LIMA-derived centrality metrics (degree, in-degree, out-degree, betweenness, eigenvector), and five using the same metrics from CellOracle-inferred GRNs. A random permutation of expressed TFs served as a negative control.

GeneSys-simulated profiles consistently outperformed unregularized scRNA-seq data across all methods (Figure 4A, 4B). DE-based strategies demonstrated the strongest and most consistent performance, surpassing network-based approaches across all data sources tested. Importantly, across all metrics examined, including tissue- and cell-type–specific analyses, GeneSys-enhanced prioritization yielded superior results, underscoring the model’s utility for regulatory inference and TF discovery (Figure 4C).

**Figure 4.**
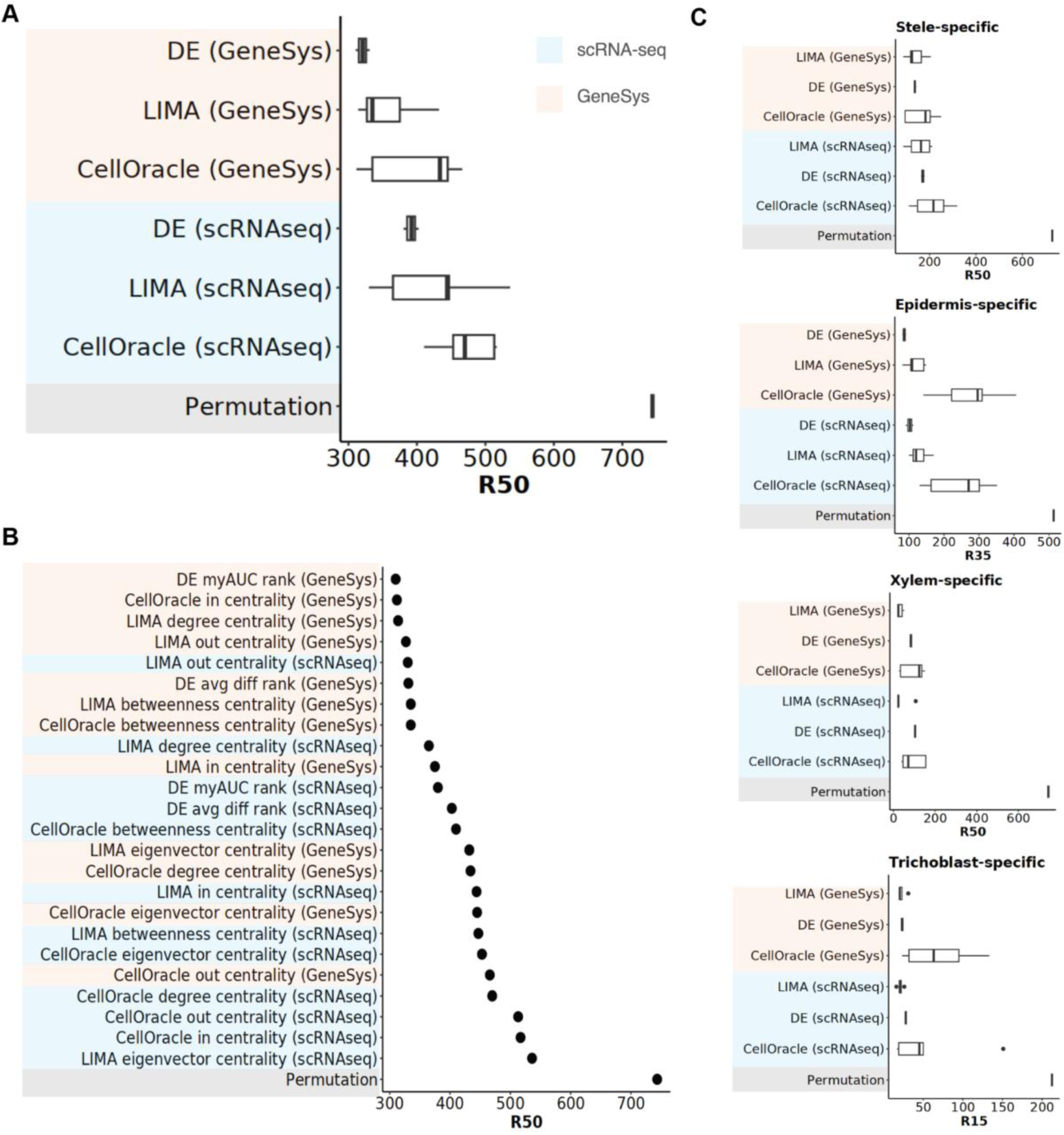
Evaluation of gene prioritization methods using a gold-standard transcription factor (TF) list associated with root development and biology. The R50 metric is defined as the rank position in a gene prioritization list at which half of the genes in the gold-standard TF set appear above that rank. Lower R50 values indicate better prioritization performance. “GeneSys” denotes prioritization schemes applied to GeneSys-simulated root transcriptomes, while “scRNA-seq” refers to schemes applied to the original root atlas data. Three gene prioritization strategies were evaluated: **DE** : based on differential expression analysis, **LIMA** : based on linear interaction matrices capturing transcriptomic changes during state transitions, **CellOracle** : based on gene regulatory networks (GRNs) inferred by CellOracle^5^. A permutation-based control was also included, calculated as the mean R50 from 1,000 random permutations of all expressed TFs. (A) Summary of benchmarking results across all methods at the system-wide level. (B) Performance comparison of individual prioritization schemes using specific ranking or centrality metrics. **DE avg diff rank** : genes ranked by the log fold-change of average expression between the target and background groups. **DE myAUC rank** : genes ranked by the area under the ROC curve (AUC), where an AUC of 1.0 indicates perfect separation of the two groups by gene expression alone. **Betweenness centrality** measures the extent to which a node lies on the shortest paths between other nodes. **Out-degree centrality** and **in-degree centrality** refer to the unweighted number of out-going and in-coming edges a node has in a network. **Degree centrality** is the sum of out-degree and in-degree centrality. **Eigenvector centrality** measures the extent to which a node that are not only well-connected but also connected to other well-connected nodes. (C) Benchmarking summary at the tissue level. (D) Benchmarking summary at the cell type level.

### Characterizing transcriptomic changes during state transitions

In addition to prioritizing transcription factors based on transcriptomic differences across tissues and cell types, GeneSys enables the characterization and comparison of subtle changes during state transitions through its temporal modeling component, TD-VAE. Leveraging the generative nature of the model, GeneSys can produce cell-by-gene expression matrices for any specified state and of any sample size, making it straightforward to derive LIMAs between any two states. These LIMAs represent the linear transcriptomic changes that coincide with a transition from one state to another. By generating LIMAs for all transitions of interest, we can comprehensively map the spatiotemporal transcriptomic dynamics of any gene within a defined developmental system.

Using the root system as an example, we identified five biologically representative and relevant state transitions (Figure S5B, see Materials and Methods): stem cell to proliferation domain (t0– t1), proliferation domain to transition domain (t1–t3), transition domain to early elongation zone (t3–t5), early to late elongation zone (t5–t7), and late elongation zone to maturation zone (t7–t9). For each transition, we derived ten LIMAs, each representing a major root cell type, resulting in a total of 50 LIMAs that together provide a comprehensive landscape of spatiotemporal state transitions in the root.

We then examined the centrality scores of genes of interest across all 50 LIMAs to depict their spatiotemporal transcriptomic dynamics. For example, we observed the well-characterized TF SHORTROOT (SHR), known for its crucial role in endodermis cell fate.^30^ The centrality scores revealed that SHR is involved in the earliest transitions within both the stele and endodermis, and continues to play roles in later transitions affecting the pericycle, xylem, and procambium (Figure 5A). These results highlight SHR’s importance in the establishment of endodermis and stele identities, consistent with findings from single-cell profiling of the *shr* mutant, where endodermis and several stele cell types are compromised^12^.

**Figure 5.**
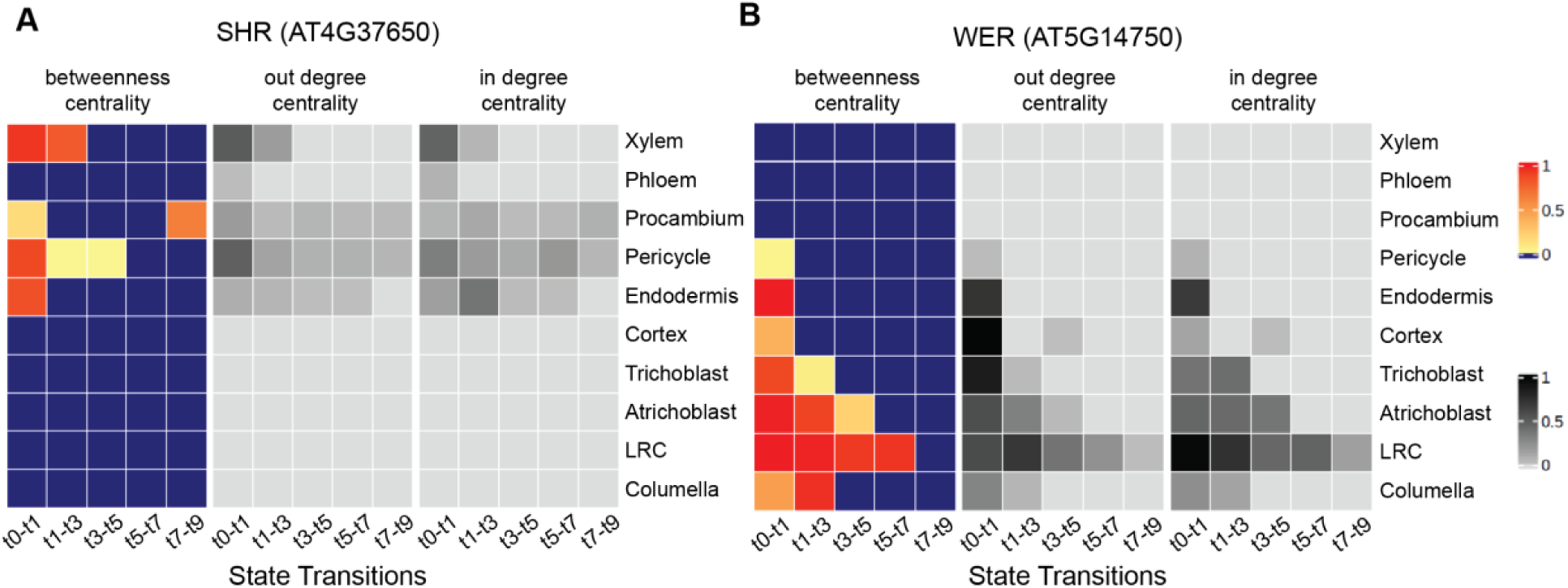
Mapping spatiotemporal gene activity during state transitions using linear interaction matrices (LIMAs) (A-B) Betweenness, out-degree, and in-degree centrality scores across all cell types and transitions for two key cell fate regulators: SHR (SHORTROOT) and WER (WEREWOLF). These transcription factors are predicted to exhibit broad, cross-tissue regulatory activity during developmental transitions.

Similarly, we analyzed WEREWOLF (WER), a TF crucial for epidermis cell fate^31^. WER’s centrality scores across LIMAs showed strong involvement in all epidermal cell types (atrichoblast, trichoblast, lateral root cap, and columella) (Figure 5B). Intriguingly, WER also appeared to participate in early transitions in the ground tissue layers (cortex and endodermis) and the outermost stele layer (pericycle), suggesting that WER may coordinate with other TFs critical to these lineages. These observations, however, should be interpreted with caution. They may reflect previously unrecognized roles of WER, but could also represent artifacts stemming from uncertainty in cell type annotations, since GeneSys depends on both the provided annotations and the cell lineage blueprint to perform data imputation. Follow-up experimental validation will be essential to confirm these predictions.

Beyond identifying TFs with cross-tissue activity, LIMA-derived centrality profiles also highlighted transcription factors with strong cell type-specific roles. To quantify specificity, we calculated the proportion of each TF’s total centrality, defined as the sum of betweenness, in-degree, and out-degree scores, contributed by each cell type across all LIMAs. A TF was classified as cell type-specific if over 50% of its total centrality was concentrated in a single cell type. Notable examples include MYB36^22^, SCR^32^, and BLJ^33^, all critical for endodermis development (Figure S10A, S10B), as well as GL2^34^ (atrichoblast), RHD6^35^ (trichoblast), JKD^36^ (cortex), APL^23^ (phloem), and VND family TFs^24^ (xylem), consistent with their established roles in root cell fate specification.

Using combined centrality scores, we also identified candidate TFs with potential roles in underexplored cell types or novel developmental functions, offering promising targets for future validation (Figure 5, Figure S10, Table S3).

In summary, by consulting known TFs using GeneSys-derived LIMAs, we gain higher resolution and deeper insights into cross-tissue transcriptomic dynamics throughout temporal developmental progression, offering a powerful framework for hypothesis generation and a deeper understanding of how gene interaction networks unfold within the spatiotemporal landscape of a developmental system.

### GeneSys simulation for the time-series data of mouse embryo development

In contrast to the relatively stable and lineage-committed structure of the *Arabidopsis* root, mouse embryogenesis presents a highly dynamic and heterogeneous developmental landscape.

To test the robustness and cross-species applicability of GeneSys, we trained the model on one of the largest time-series scRNA-seq datasets to date^13^, comprising 11.4 million nuclei from 74 embryos spanning embryonic day 8 (E8) to postnatal day 0 (P0). Snapshots were collected at 2-6 hour intervals, resulting in 43 temporally resolved stages.

To define major developmental windows, we performed hierarchical clustering of pseudobulk transcriptomes across all time points, identifying eleven distinct developmental stage clusters. After excluding transient (e.g., primitive erythroid) and extremely rare (e.g., testis, adrenal) cell types, we focused on twenty-four major cell types. Together with the eleven stage clusters, these defined the cell type trajectories used for GeneSys training (Figure S3B, S5D).

Despite the large sample size, the data were considerably sparser than the *Arabidopsis* atlas. After filtering low-quality nuclei (< 2500 genes or UMI counts), we retained ∼1.5 million high-quality profiles. We further downsampled to a maximum of 500 nuclei per cell type–stage combination, yielding a balanced set of ∼100 k cells. As with the *Arabidopsis* dataset, data were split into training (64%), validation (16%), and test (20%) sets. Each cell expressed 24552 genes, and training batches were assembled into a tensor of shape (24, 11, 24552), sampled based on a predefined cell lineage blueprint.

Unlike in roots, many mouse cell types emerge later in development, often without clearly annotated precursors. In such cases, e.g., definitive erythroid cells or white blood cells, early stages were represented by zero-filled expression vectors. The lineage blueprint (Figure S3B) encoded relationships such as neuroectoderm giving rise to neurons and glia, mesoderm to muscle and adipocytes, and epithelial cells to lung and airway lineages.

Following training, GeneSys-generated transcriptomes exhibited greater continuity than the original test data. While real test batches often contained sparse and disjointed profiles, the model-generated trajectories were regularized and interconnected (Figure 6). Critically, the simulated developmental paths recapitulated known cell type-specific marker gene expression patterns^13^. For example, Pax1 (mesoderm), Gad1 (CNS neurons), Cspg4 (oligodendrocytes), Myf5 (muscle cells), and Alb (hepatocytes) were all robustly expressed in the appropriate model-derived lineages, consistent with established developmental markers in the mouse embryo (Figure S11).

**Figure 6.**
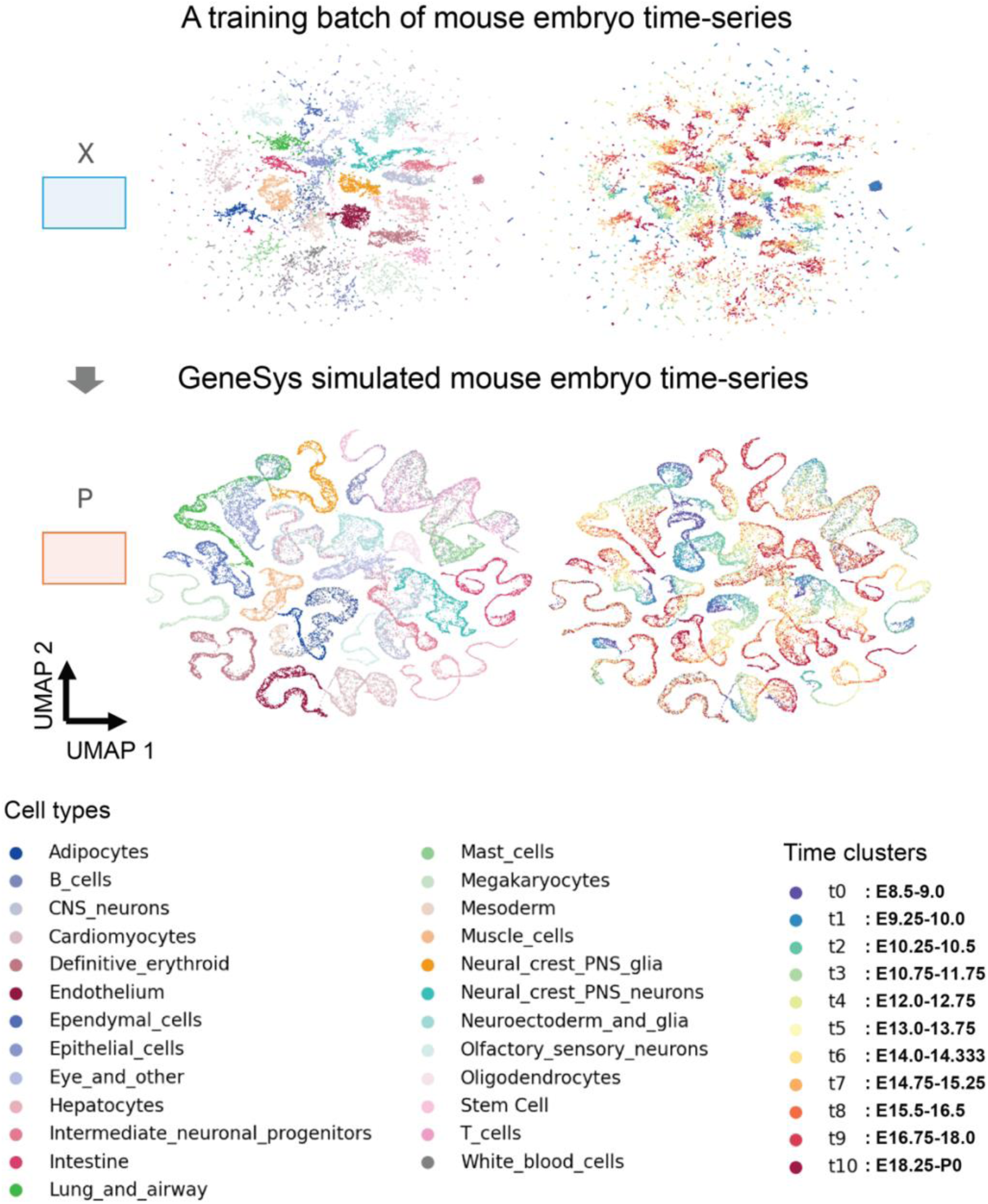
The trained GeneSys model accurately simulates the mouse embryo development time-series and generates dense, regularized single-cell transcriptomes. UMAP embeddings of a representative training batch X, and the GeneSys-simulated mouse embryo development time-series P. The training batch X and the simulated data P contain the same number of cells. (B) UMAPs showing the expression of known marker genes in the original and simulated mouse embryo development time-series, highlighting the preservation of cell type-specific expression patterns.

Recreation accuracy based on masked test inputs revealed that GeneSys requires input from at least 6-7 time points to achieve satisfactory accuracy (≥0.8) (Figure S5C). This reflects the inherent complexity of mouse embryogenesis, where many cell types are absent in early stages and only differentiate later. At earlier time points, the lack of definitive lineage identity limits the model’s ability to accurately classify or predict future states. The requirement for multiple temporal inputs underscores the intricate and dynamic nature of mammalian development and highlights the capacity of GeneSys to capture gradual cell fate transitions.

Despite the contrasting complexity, GeneSys successfully reconstructed developmental trajectories in both *Arabidopsis* root and mouse embryo, demonstrating its generalizability across diverse biological systems.

## Discussion

We introduce GeneSys, a generative model designed to simulate single-cell transcriptomic landscapes in developmental systems. Its primary goal is to create a noise-reduced and robust virtual biological system by incorporating prior knowledge of cell lineage blueprints, providing an *in-silico* platform for hypothesis generation and resource prioritization.

Unlike unsupervised autoencoder models that focus on single-cell data denoising and imputation^7, 8^, GeneSys emphasizes temporal modeling while encoding domain knowledge of developmental constraints as prior knowledge. By learning representative transcriptomic dynamics along developmental trajectories, GeneSys captures both cell type specification and maturation processes, critical for understanding tissue formation and organismal development. Because GeneSys incorporates external information, specifically, biological structure in the form of cell lineage blueprints, during model training, it also helps mitigate the circularity issues often associated with imputation-based models, which can artificially inflate gene to cell correlations and introduce false-positive in downstream analysis^9^.

Applications in *Arabidopsis* and mouse embryos demonstrate the ability of GeneSys to reconstruct robust developmental trajectories, generate biologically meaningful profiles, and facilitate gene discovery. Through gene masking and augmentation, we show that GEPs encode core functional and developmental logic and can serve as interpretable units for downstream analysis.

While GeneSys does not directly identify regulatory drivers of GEP activity, it enables prioritization of candidate regulators, particularly TFs within relevant GEPs, for experimental validation (Table S1). Once key TFs are identified, their interactions across multiple GEPs can be systematically investigated to uncover higher-order regulatory coordination.

To support such exploration, the trained GeneSys model functions as a flexible platform for testing combinations of GEP perturbations. By simulating co-activation or inhibition of GEPs, GeneSys allows researchers to generate and refine hypotheses, accelerating the discovery of developmental regulatory mechanisms.

As a temporal model, GeneSys supports detailed analysis of state transitions. It can generate representative cell-by-gene expression matrices for any developmental state and sample size. LIMAs derived from paired expression matrices approximate gene-gene relationships during state transitions, enabling the construction of dynamic gene networks. These networks can be analyzed using graph-based centrality metrics to identify regulatory hubs, providing spatiotemporal insight into transcriptomic shifts.

With the rapid expansion of single-cell datasets, there is growing demand for computational frameworks and technique that can integrate high-dimensional, heterogeneous data and uncover both associative and causal relationships. Future extensions of GeneSys could incorporate multi-omic modalities (e.g., ATAC-seq, proteomics) and simulate treatment responses through advanced network modeling. As a neural network-based generative model, GeneSys is inherently modular and extensible. Its core strategy, learning latent structure from high-resolution data and interpreting developmental logic via derived gene networks, positions it as a powerful tool for studying developmental systems. GeneSys can be applied to diverse systems and schemes with annotated cell types and time points, for example, time series dataset with multiple mutants and conditions, making it broadly useful for elucidating gene interaction dynamics underlying cell fate decisions, developmental progression and responses to biotic and abiotic stimuli.

## Supplementary Figures

**Figure S1.**
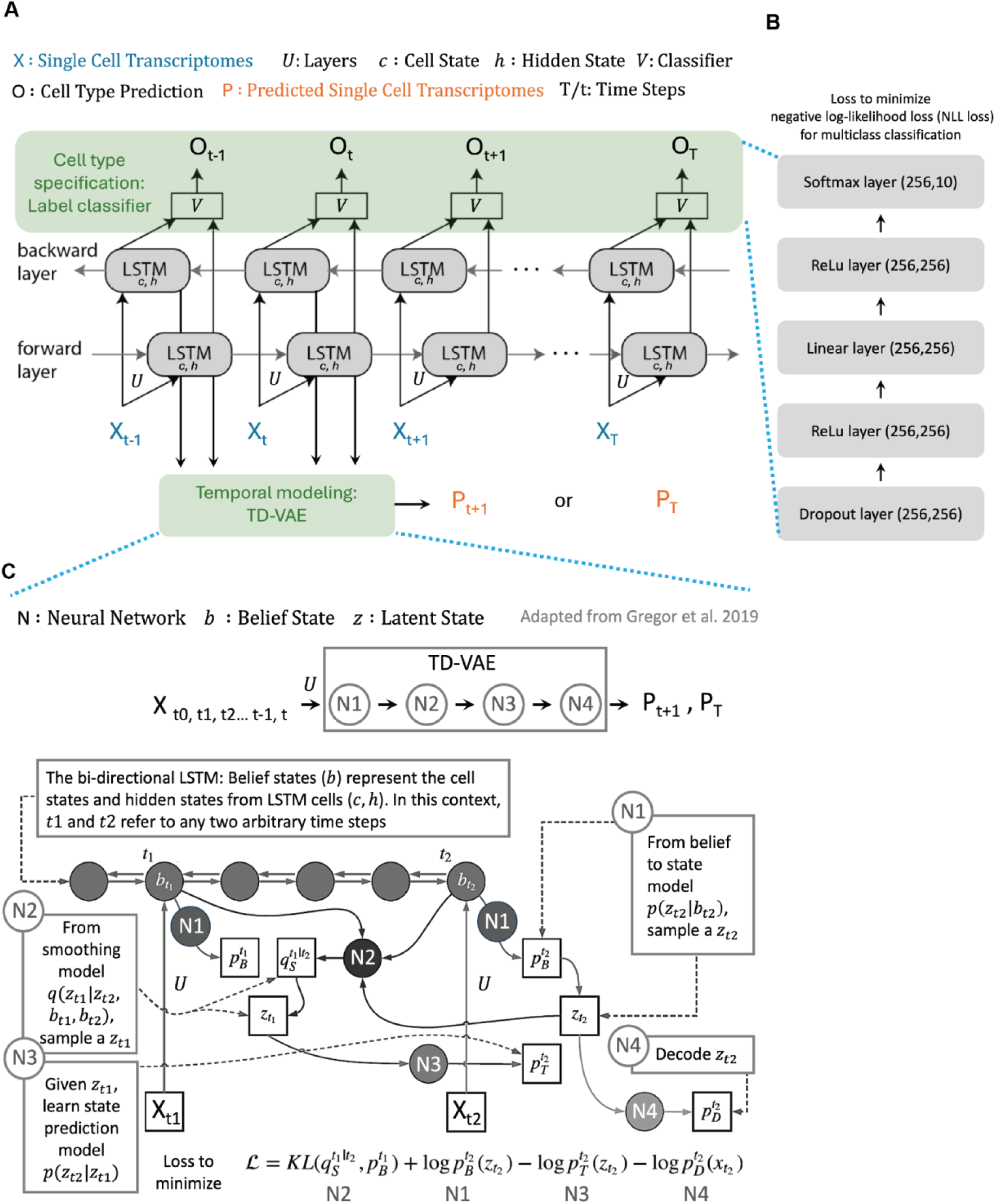
Detailed GeneSys model architecture. (A) Overview of the GeneSys architecture. A bi-directional LSTM network is connected to both a multi-label cell type classifier and a TD-VAE, enabling the model to learn cell type specificity and temporal developmental dynamics, respectively. Notation: **X** - single-cell transcriptomes; **U** - neural network layers; **c** - LSTM cell state; **h** - LSTM hidden state; **V** - cell type classifier; **O** - predicted cell type labels; **P** - predicted single-cell transcriptomes; **t** - specific LSTM time step; **T** - any LSTM time step. (B) Layer configuration of the multi-label cell type classifier. Each layer is annotated with its input and output dimensions in the format (input, output). (C) Architecture of the TD-VAE^15^. **N** - neural network components; **b** - belief state, computed as the concatenation of LSTM cell state and hidden state; **z** - latent state; **KL** – Kullback-Leibler divergence.

**Figure S2.**
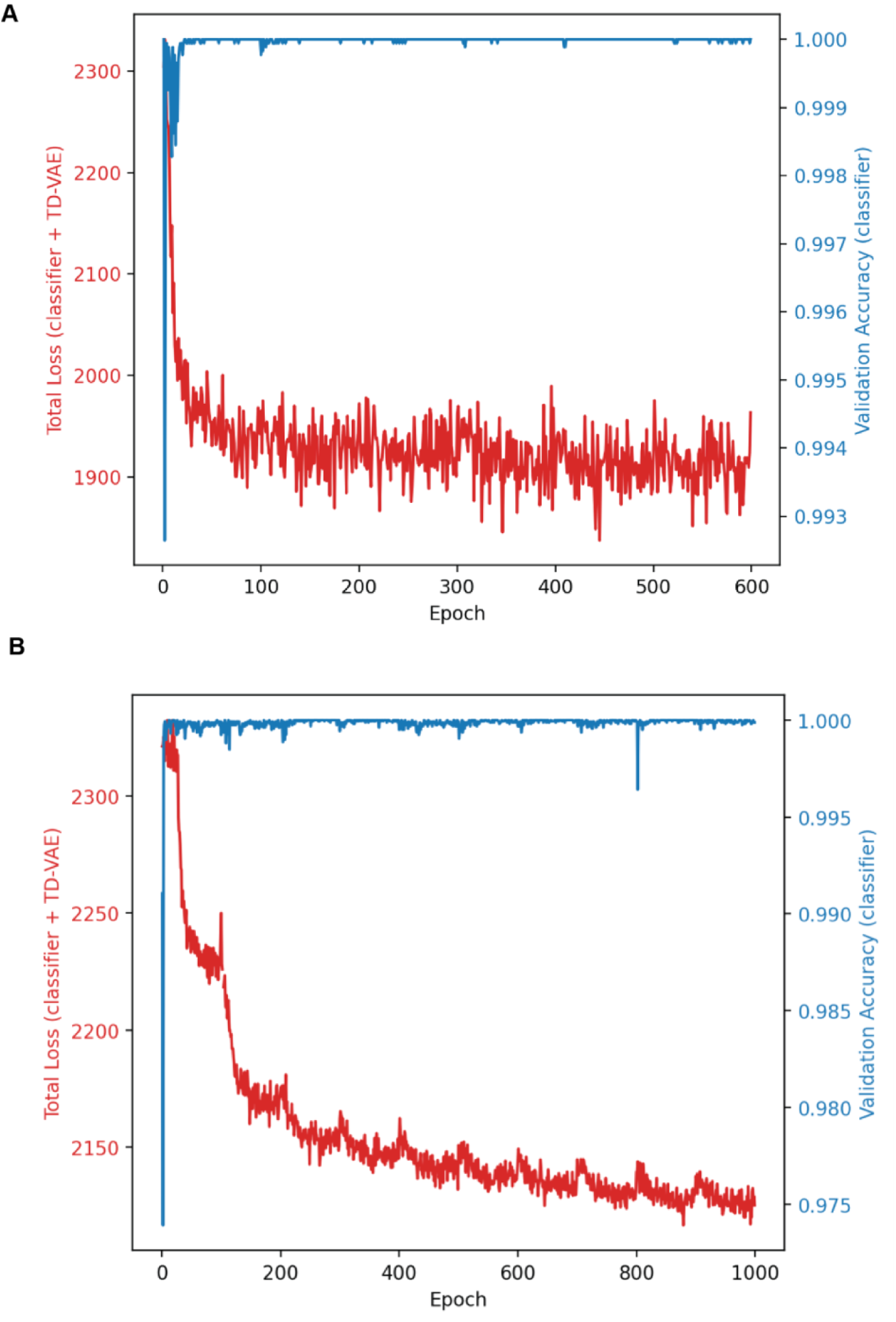
Training logs of GeneSys models. Total training loss and validation accuracy recorded across epochs for (A) the root atlas and (B) the mouse embryo time-series datasets.

**Figure S3.**
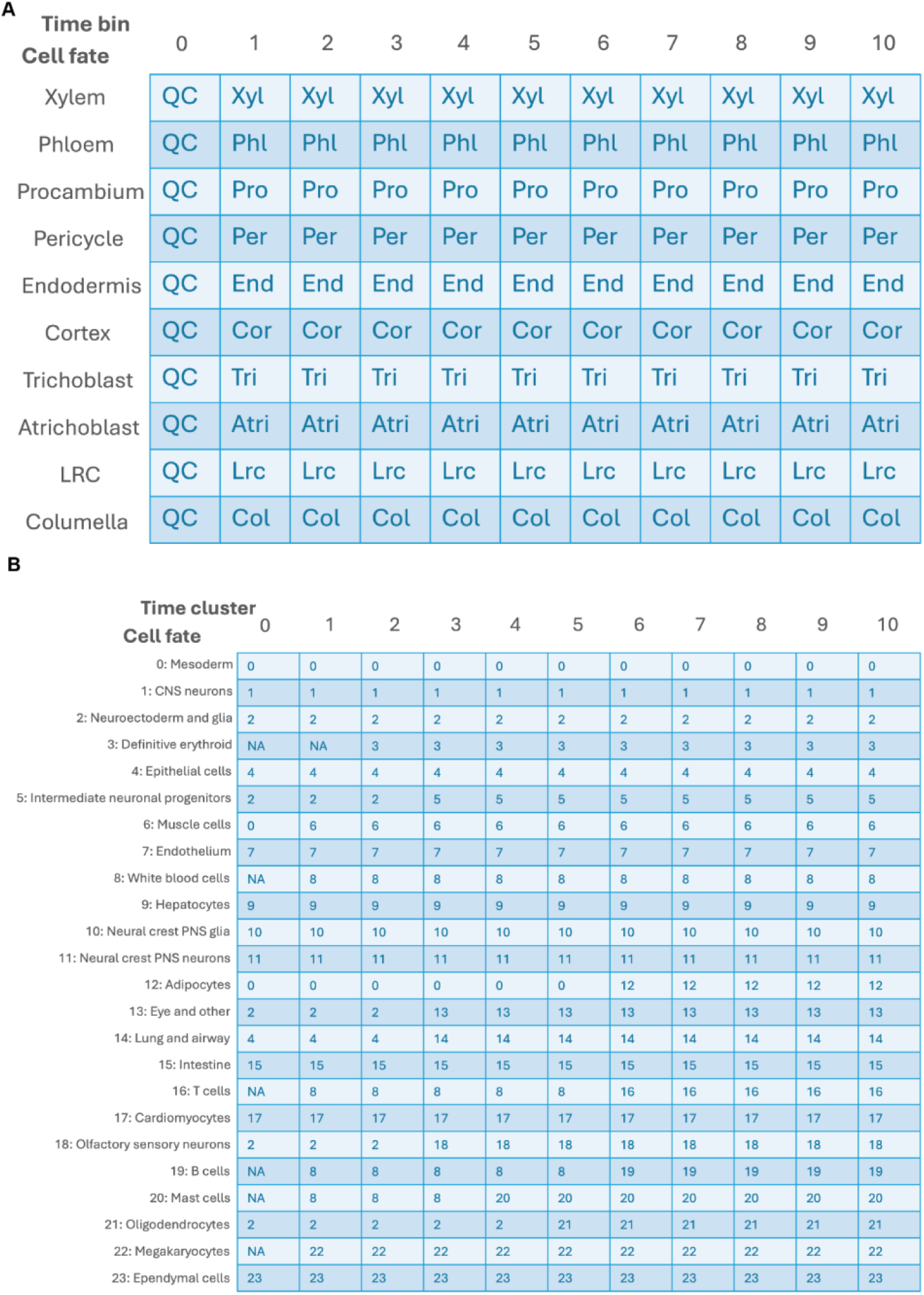
Examples of cell lineage blueprints. Cell lineage blueprints used for (A) the root atlas and (B) the mouse embryo time-series. Each tile indicates the specific cell type transcriptomes from which the model samples during training. Columns correspond to time bins (root) or developmental stage clusters (mouse). Tiles labeled “NA” represent non-existent cell states at that stage, for which zero-filled profiles were used in place of transcriptomic data.

**Figure S4.**
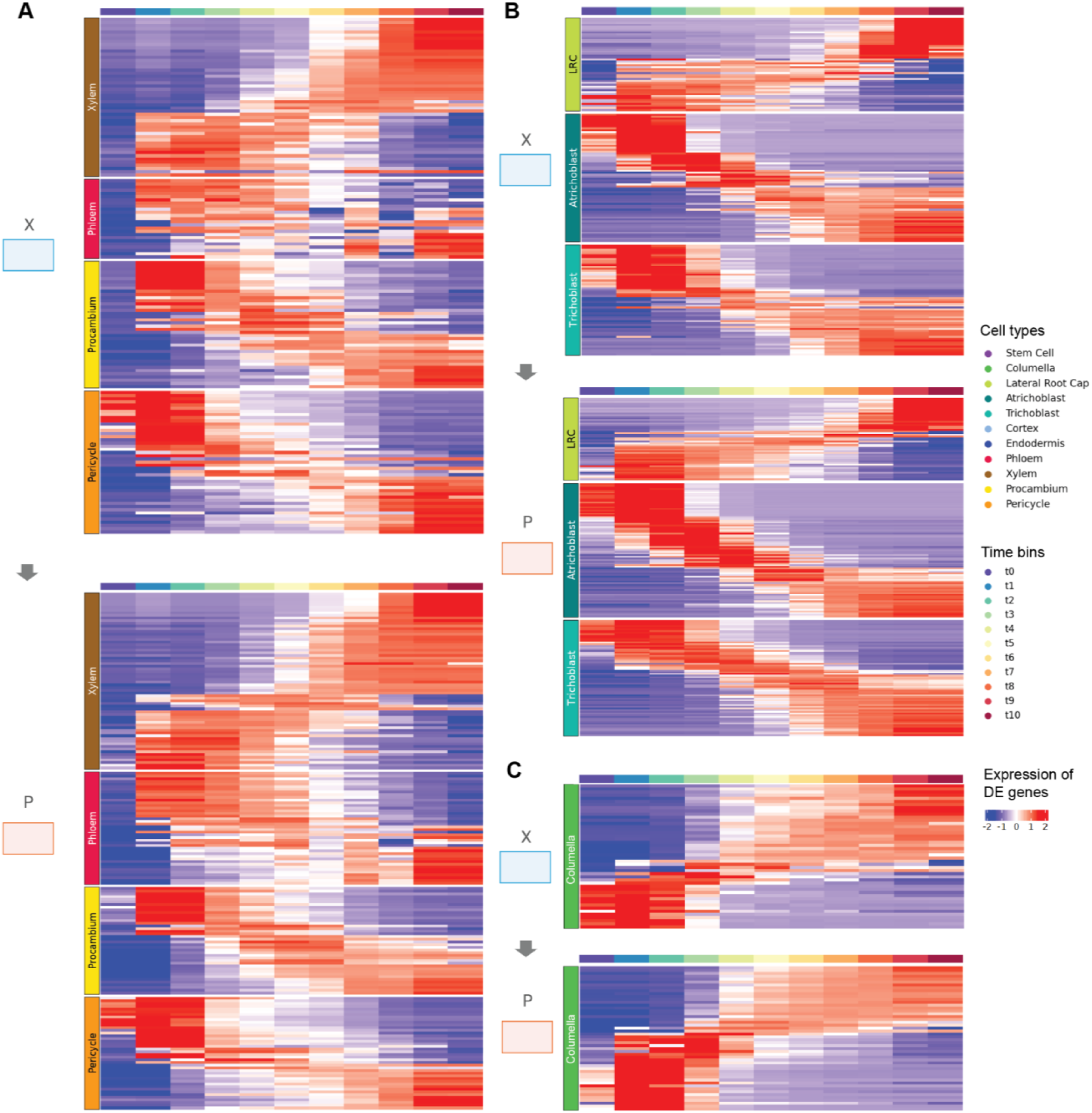
Heatmaps of differentially expressed (DE) genes along developmental trajectories in the root atlas and GeneSys-simulated data. Scaled expression of non-redundant DE genes across eleven pseudotime bins for (A) stele, (B) epidermal tissue, and (C) columella. Warmer colors indicate higher expression levels. Although thousands of DE genes were identified across pseudotime, only the most strongly differentially expressed genes for each of the eleven pseudotime bins are shown for clarity.

**Figure S5.**
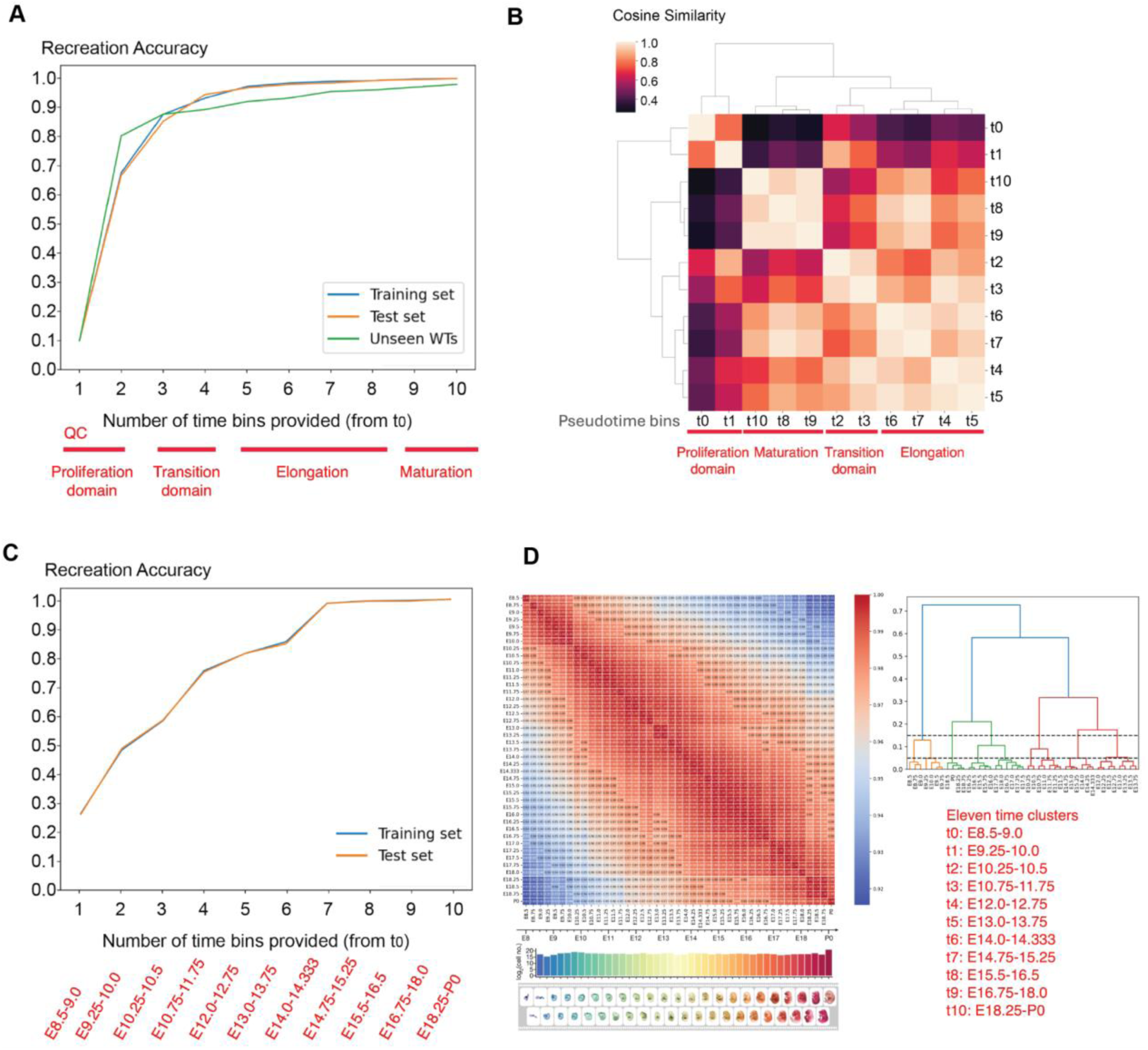
Recreation accuracy and temporal annotations of developmental time bins/clusters. (A, C) Recreation accuracy of GeneSys in the *Arabidopsis* root (A) and mouse embryo (C) time-series datasets, based on progressively increasing numbers of preceding time bins provided as input (x-axis, starting from t₀). Recreation accuracy (y-axis) quantifies the model’s ability to reconstruct developmental trajectories from partial input. (B) Cosine similarity among transcriptomes across time points in the root atlas. Five clusters were identified, corresponding to the proliferation domain (t₀–t₁), transition domain (t₂–t₃), early elongation zone (t₄–t₅), late elongation zone (t₆–t₇), and maturation zone (t₈–t₁₀). (D) Cosine similarity among transcriptomes across developmental stages in the mouse embryo time-series. Hierarchical clustering revealed eleven distinct stage clusters.

**Figure S6.**
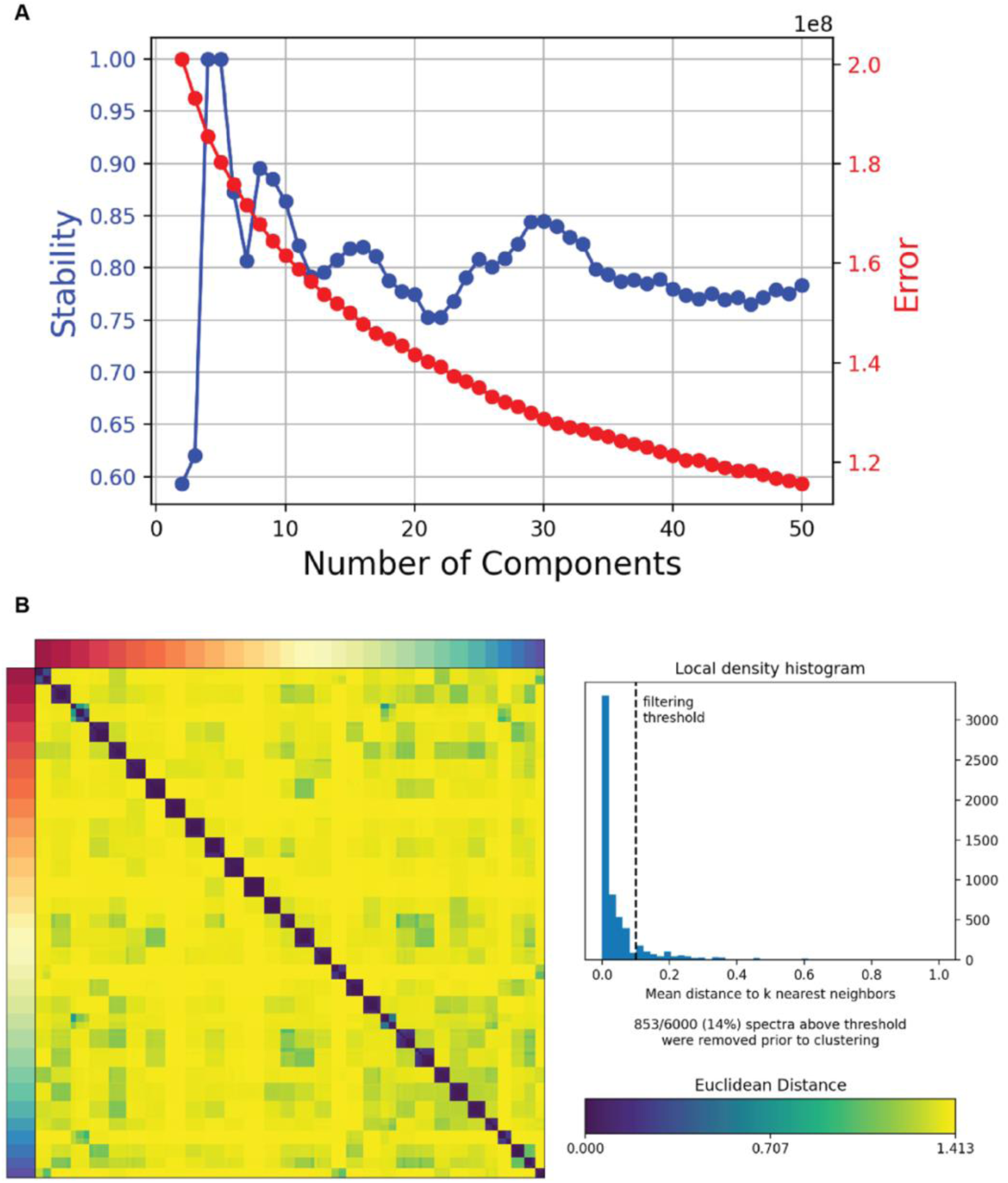
Gene expression programs (GEPs) identified in the WT root atlas^12^. Thirty GEPs were identified in the root atlas (Shahan and Hsu et al., 2022)^12^ using consensus non-negative matrix factorization (cNMF). After matrix factorization, the cell-by-gene matrix is decomposed into two matrices: a cell-by-factor matrix and a factor-by-gene matrix. Here, each factor corresponds to an identified GEP. The factor-by-gene matrix captures the relationship between each gene and each GEP, while the cell-by-factor matrix contains the loadings/weights of each GEP in individual cells. In this work, we refer to these loadings as GEP usage. (A) Scanning for optimal K with the highest stability beyond tissue/cell type level information (K <= 10). (B) To refine gene assignments, Euclidean distances among genes were computed, and genes with a mean distance to their k nearest neighbors below 0.1 were excluded.

**Figure S7.**
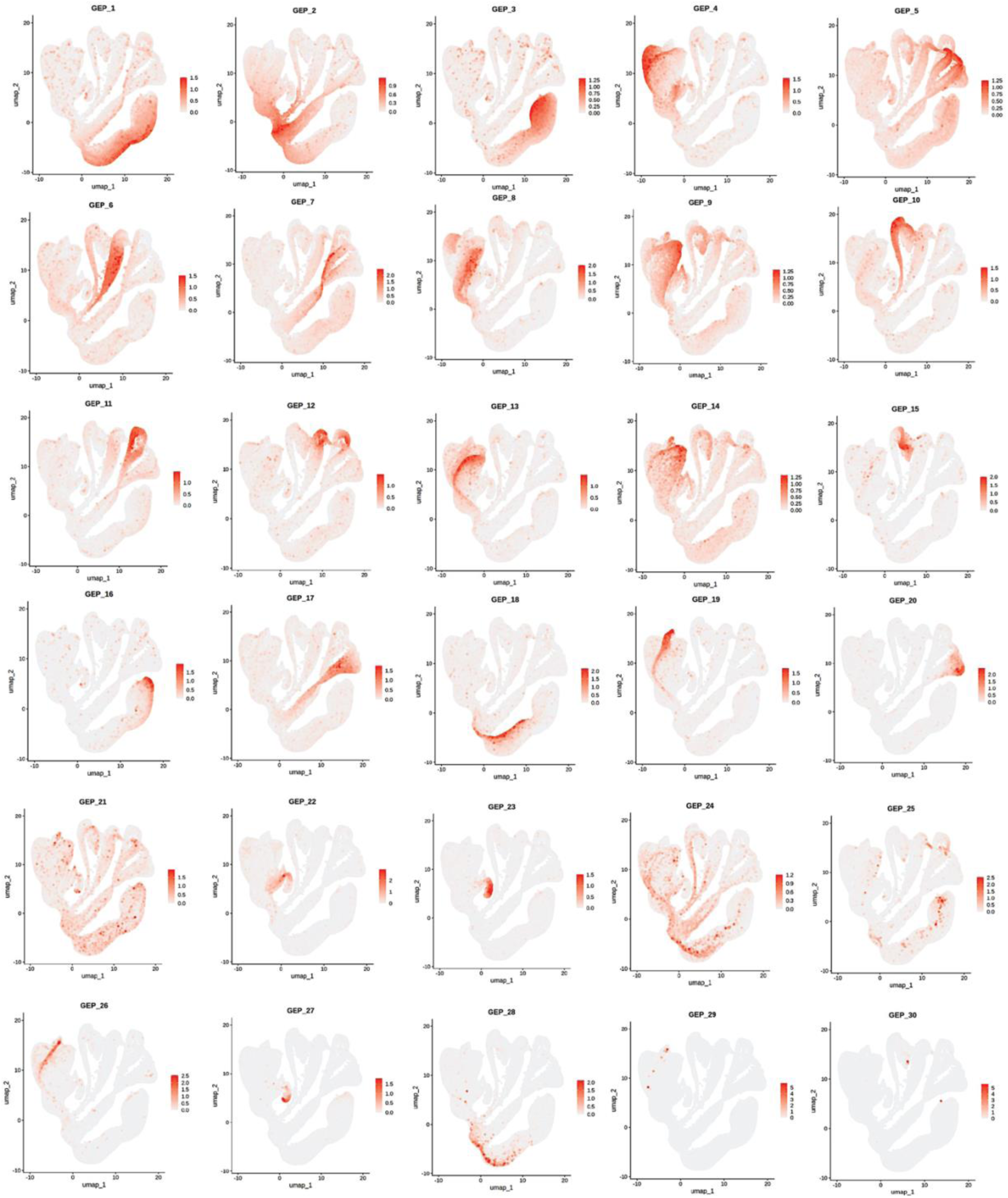
Gene expression programs (GEPs) usage on the UMAP of the root atlas^12^.

**Figure S8.**
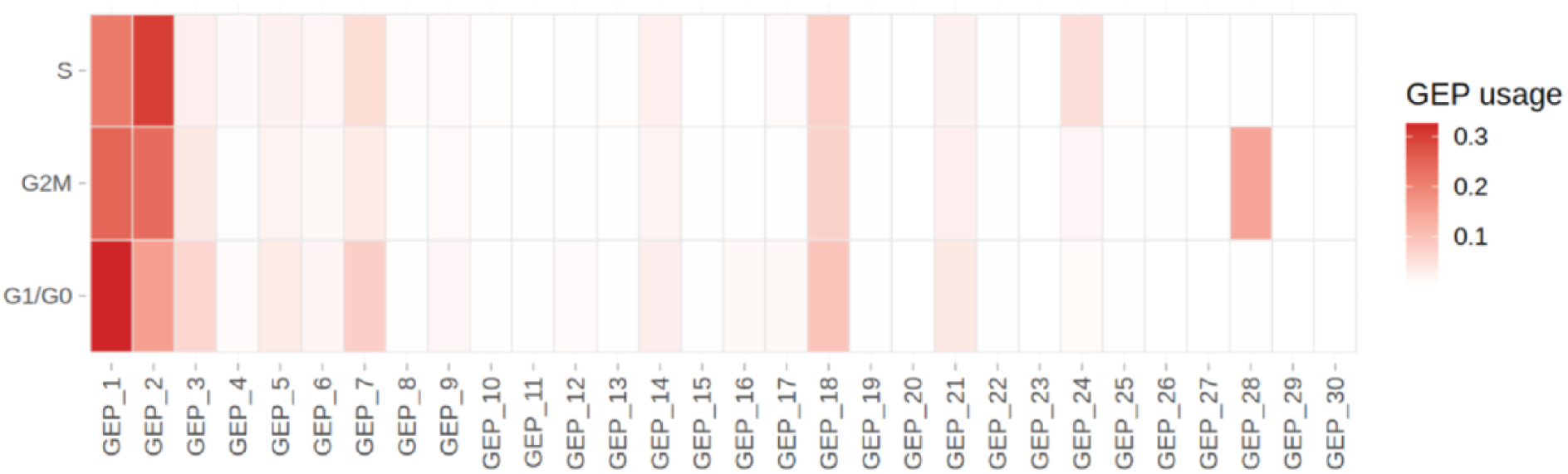
GEP averaged usage across annotated cell cycle phases^37^. Only “Proliferation Domain”, “Proximal Lateral Root Cap”, “Proximal Columella” cells where active cell cycle is present are included for the calculation

**Figure S9.**
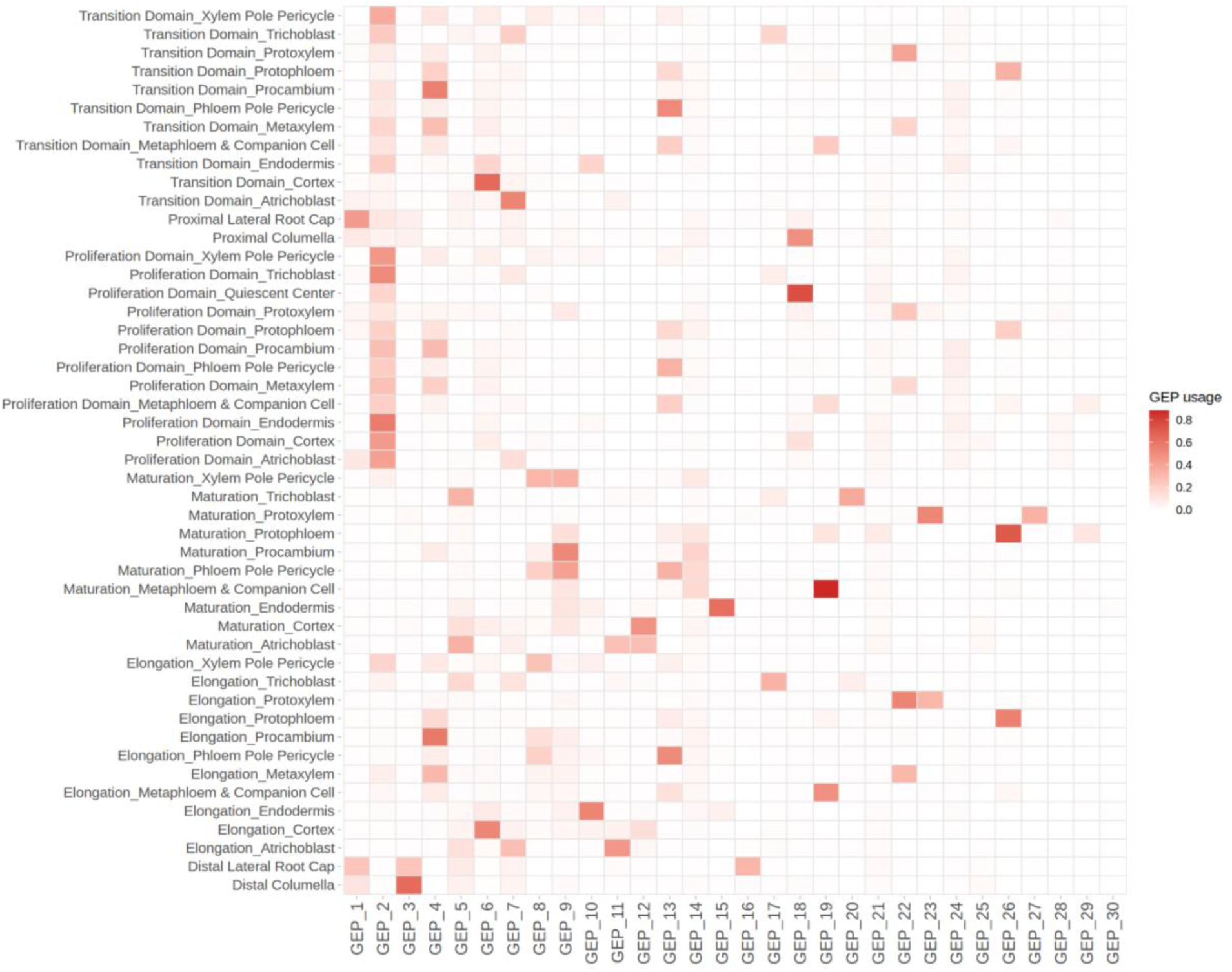
Gene expression programs (GEPs) averaged usage across annotated cell types, developmental stages^12, 25^.

**Figure S10.**
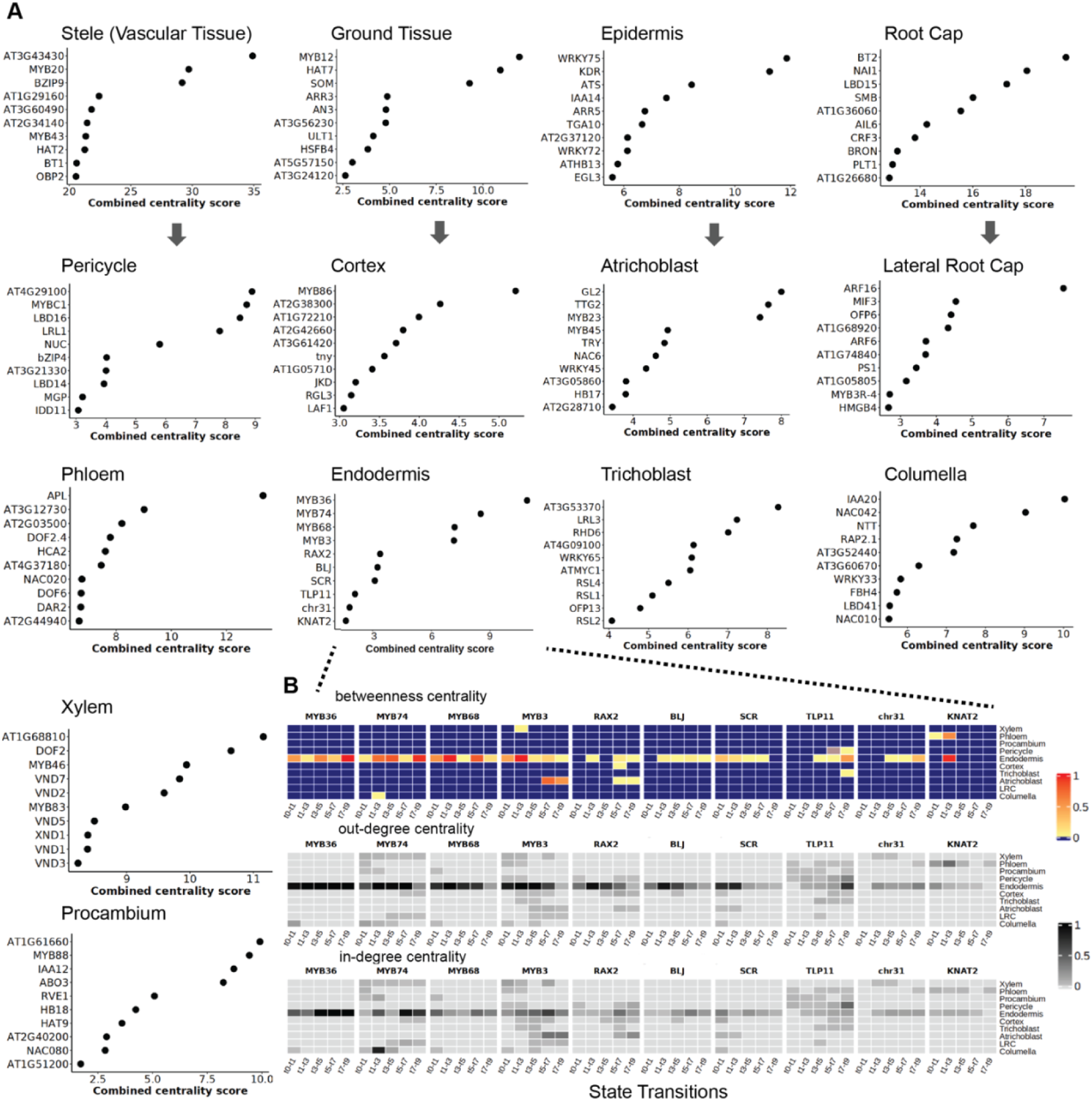
Ranking gene importance during spatiotemporal state transitions with linear interaction gene networks and centrality measurements. (A-B) Transcription factors were ranked using a combined centrality score, calculated as the sum of betweenness, out-degree, and in-degree centrality across all transitions, based on GeneSys-derived LIMAs. (A) Rankings of transcription factors at both tissue and cell type levels. (B) Centrality profiles across all transitions for top-ranked transcription factors associated with endodermis fate, shown as an example.

**Figure S11.**
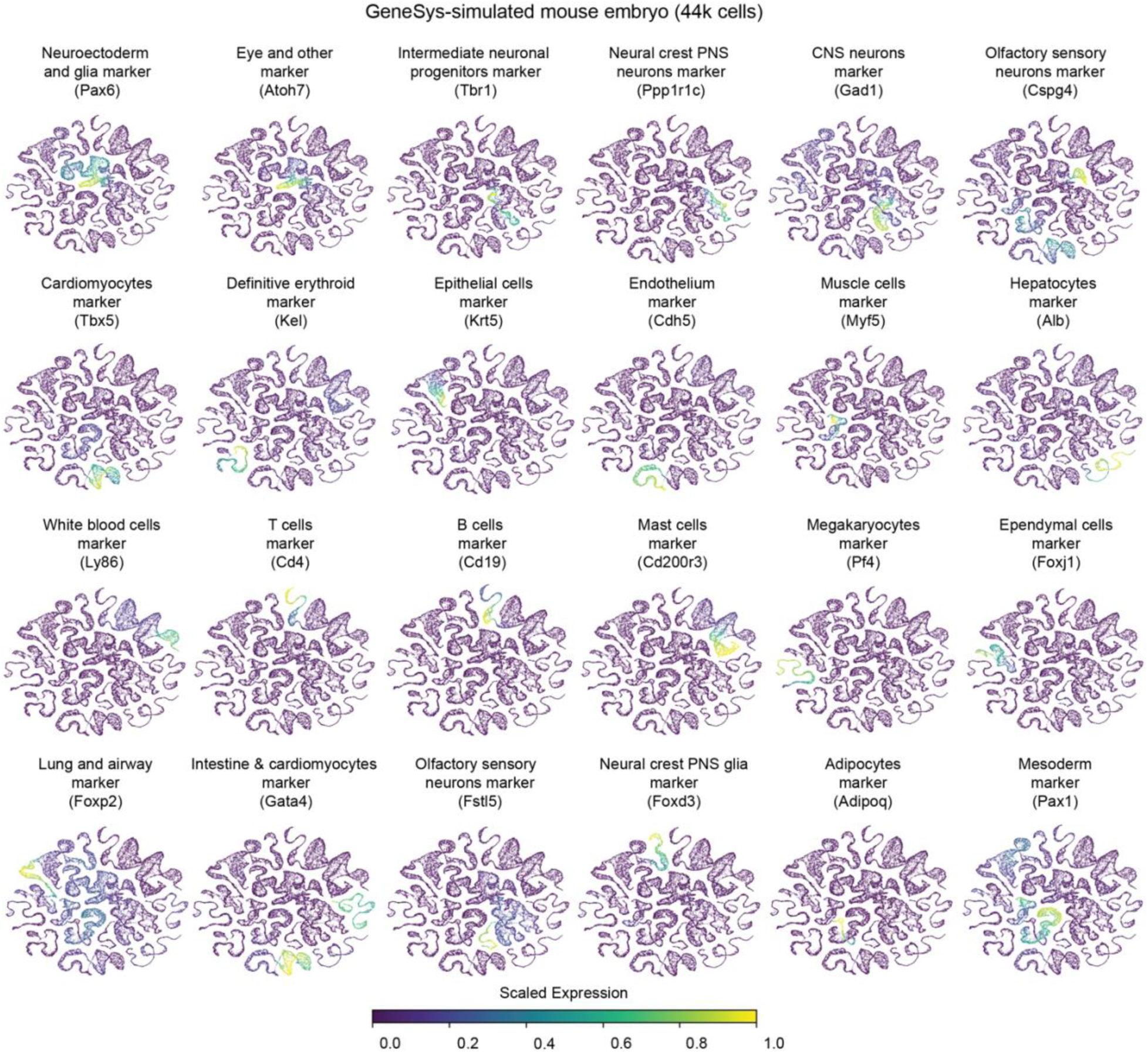
Known marker gene expression patterns on GeneSys-simulated mouse embryo time-series^13^.

## Materials and Methods

### Lineage-blueprint-guided trajectory construction and training strategy for GeneSys

To model temporal dynamics across developmental lineages, training batches were constructed based on a predefined cell lineage blueprint. For a hypothetical system comprising three cell types, A, B, and C, progressing through six developmental stages, each training batch was formatted as a tensor of shape (3, 6, n), where n is the number of genes profiled. For each cell type, transcriptomes were sampled according to their lineage-defined origin. For example, the trajectory of cell type C was constructed by sampling transcriptomes from cell type A at stages 1 and 2, cell type B at stages 3 and 4, and cell type C at stages 5 and 6, reflecting the progression from A to B to C across development. The same approach was applied to cell types A and B using their respective lineage paths. These assembled trajectories were used as input to the GeneSys model (Figure 1B).

### Root data preparation for GeneSys training and evaluation

Seurat^38–40^ v4.1.0 objects for the root atlas and individual samples were downloaded from GEO (accession: GSE152766). The input for GeneSys consisted of single-cell transcriptomes, cell type annotations, and inferred pseudotime values for each annotated cell. The Seurat objects (.rds) were converted into Scanpy-compatible AnnData objects (.h5ad) using Seurat (v4.1.1.9001) and SeuratDisk (v0.0.0.9020)^39^.

For model input, batch-corrected and scaled expression values from the integrated assay of the Seurat objects were used as the primary representation of single-cell transcriptomes. Separately, scaled SCTransform-normalized values (SCT assay) were also evaluated and shown to produce comparable results when used for training. Expression values were floored at zero and normalized to the [0, 1] range. The resulting normalized cell-by-gene matrices were split into three subsets: Training set (65%) used for model training; Validation set (15%) used for parameter tuning during training; Test set (20%) used to evaluate the performance of the model, including the accuracy of the cell type classifier on generated single-cell transcriptomes.

Annotations for eleven major cell types: Quiescent Center, Endodermis, Cortex, Atrichoblast, Trichoblast, Xylem, Phloem, Pericycle, Procambium, Columella, and Lateral Root Cap, were extracted directly from the Seurat objects. The Quiescent Center (QC) was designated as the root system’s stem cell population and assigned to time bin zero. For the remaining cell types, pseudotime trajectories were divided into ten equal-sized time bins using the ‘consensus pseudotime’ values provided in the atlas and the qcut function from the pandas package (v1.2.4). For evaluation on unseen data, i.e., samples not used during training, annotations for wild-type samples sc_20 and sc_21 (GSE152766) were transferred from the root atlas following Seurat’s label transfer workflow^38, 40^. Pseudotime inference for these samples was performed using CytoTRACE v0.1.0^41^, and binning was applied as described above.

Each cell in the root atlas was profiled for 17513 genes. To construct model inputs, developmental trajectories were represented as matrices of shape (11, 17513), where the eleven time steps correspond to pseudotime bins along the differentiation and maturation axis. Given the presence of ten major cell types in the atlas (excluding quiescent center, QC), an initial batch was defined as one trajectory per cell type, resulting in a tensor of shape (10, 11, 17513).

During training, single-cell transcriptomes were sampled according to a predefined cell lineage blueprint (Figure S3A), ensuring balanced representation across both cell types and pseudotime bins. To assemble a training batch, trajectories were randomly sampled from the training set, with each trajectory replicated approximately 51 times to match a predefined batch size of 512. This yielded a final training batch of shape (512, 11, 17513): 512 trajectories per batch, each spanning 11 pseudotime steps and 17513 gene expression features.

The GeneSys model employed a Long Short-Term Memory (LSTM) network, operating over sequences of length 11, with each LSTM cell corresponding to a specific pseudotime bin. The hidden states generated by the LSTM were passed to two downstream components: a multi-label classifier to learn cell type specificity, and a temporal difference variational autoencoder (TD-VAE) to capture maturation dynamics across pseudotime (Figure 1B).

Detailed code for preparing the root data and executing GeneSys training and evaluation is provided in Jupyter Notebooks 0-4, available via the GitHub repository (see Code Availability section).

### Mouse embryo data preparation for GeneSys training and evaluation

Mouse embryo time-series dataset was downloaded from https://omg.gs.washington.edu/jax/public/download.html, along with the accompanying cell type annotations. Cells with fewer than 2500 unique molecular identifier (UMI) counts or fewer than 2500 detected genes were filtered out to ensure data quality.

To define key developmental windows, we computed pairwise correlations among pseudobulk transcriptomes across all 43 developmental stages and performed hierarchical clustering. This analysis identified 11 distinct time clusters, representing major stages of mouse embryonic development. We excluded primitive erythroid cells (a transient early cell type), as well as testis and adrenal cells (rare populations), resulting in a curated set of 24 major cell types. These 24 cell types, in combination with the 11 time clusters, define the cell type trajectories used for training GeneSys (see Figure S3B and Figure S5D).

To balance the dataset and reduce computational cost, we downsampled the data so that each unique state (i.e., a specific cell type and time cluster combination) contained a maximum of 500 cells. States with fewer than 20 cells were excluded and treated as missing. Expression values were scaled and floored at zero and normalized to the [0, 1] range. The resulting normalized cell-by-gene expression matrices were then divided into three subsets: Training set (65%) used for model training; Validation set (15%) used for parameter tuning during training; Test set (20%) used to evaluate model performance, including cell type classification accuracy on simulated single-cell transcriptomes.

During training, single-cell transcriptomes were sampled according to a predefined cell lineage blueprint (Figure S3B) to form input training batches. Each batch was assembled into a tensor of dimensions (512, 11, 24552): 512 cells sampled across the 24 major cell types; 11 time steps representing developmental stage clusters; 24552 genes used for training. For cell state with missing data, cells were sampled from adjacent states for imputation.

Detailed code for preparing the mouse embryo dataset and executing GeneSys training and evaluation is provided in Jupyter Notebook 5-6, available via the GitHub repository (see Code Availability section).

### GeneSys model

The GeneSys model was formulated, trained, validated, and tested using functions from the Python package PyTorch (v1.13.0). Each input, representing a single-cell transcriptome, had a dimensionality equal to the number of genes/features in the system (denoted X in Figure 1A). These inputs were first passed through an embedding layer (U in Figure 1A), consisting of the following components: A linear layer: (number of genes/features, 256), A dropout layer with a dropout probability of 0.2: (256, 256), and a Gaussian noise regularizer with σ = 0.2: (256, 256). The resulting 256-dimensional embeddings were then fed into a bi-directional LSTM network with hidden layers of size 256 and a dropout probability of 0.2. The hidden states of each LSTM cell (h in Figure 1A) were branched into two parallel paths: one for a multi-label cell type classifier, and the other for a temporal difference variational autoencoder (TD-VAE).

Within the LSTM network, the cell state functions as a memory bank, traversing the LSTM cell chain while selectively retaining or discarding information through internal gating mechanisms. In contrast, the hidden state, derived from the cell state, contains information deemed pertinent for passing to the next time step. It undergoes further processing, including activation functions and output gating, before being output from the LSTM cell for subsequent tasks or propagation to downstream layers (classifier and TD-VAE) in the neural network.

The multi-label classifier (V in Figure 1A) had the following architecture (Figure S1B): A dropout layer (p = 0.2): (256, 256), A ReLU activation layer: (256, 256), Two linear layers, each followed by ReLU layer: (256, 256), A final linear layer: (256, number of cell types/fates), and a Softmax activation layer to output predicted probabilities for each cell type (O in Figure 1A).

The TD-VAE followed the architecture described in Gregor et al. (2019) (Figure S1C), with all latent layers having input and output dimensions of 256. The decoder of the TD-VAE had dimensions (256, number of genes/features) and was used to generate predicted single-cell transcriptomes for the next time steps (P in Figure 1A).

Model training used the AdamW optimizer^42^ with an initial learning rate of 0.001. A learning rate scheduler (ReduceLROnPlateau) was implemented with parameters factor = 0.5, patience = 10, threshold = 0.05 to decrease the learning rate when improvement in the GeneSys loss plateaued. The overall GeneSys loss combined the negative log-likelihood (NLL) loss from the classifier and the TD-VAE loss.

During each training epoch, a random time bin (between t₁ and t₉) was selected for loss computation and backpropagation. Either the NLL loss or the TD-VAE loss was randomly chosen to update network weights. For every 100 epochs, the learning rate was reset to 0.001 to prevent premature convergence and potential local minimum trap.

The model was trained using a 32 GB Tesla V100 SXM2 GPU until convergence was achieved, defined as no further decrease in total loss over a rolling window of 100 epochs. The model weights corresponding to the lowest total loss across all epochs were saved and used for downstream evaluation.

Detailed instructions and code for configuring and training the GeneSys model are provided in Jupyter Notebook 1-3 and 5-6, available in the GitHub repository (see Code Availability section).

### Generation and evaluation of simulated developmental trajectories

The trained GeneSys model was used to generate simulated developmental trajectories for all cell types. Input tensors were constructed by sampling transcriptomes from the test set, and the batch size, the first dimension of the input tensor, determined the total number of simulated single-cell transcriptomic trajectories generated. The output transcriptomes were processed and visualized using Scanpy v1.9.1^43^. The processing pipeline included the following steps: Scaling: scanpy.pp.scale with max_value = 10. Principal Component Analysis (PCA): scanpy.tl.pca with svd_solver = ‘arpack’. Neighborhood graph construction: scanpy.pp.neighbors with n_neighbors = 30. Clustering: scanpy.tl.leiden with default parameters. Coarse-grained connectivity mapping: scanpy.tl.paga with default parameters. Dimensionality reduction: scanpy.tl.umap with init_pos = ‘paga’. Following preprocessing, marker gene expression was visualized using scanpy.pl.umap.

### Recreation accuracy

Recreation accuracy is a metric used to evaluate the model’s ability to reconstruct developmental trajectories. It quantifies the proportion of correct predictions made by the cell type classifier on the generated single-cell transcriptomes, given input from earlier time steps. For example, if the model is provided with transcriptomes from cell type A at time bins 0 and 1, with masked time bins of the rest, it is expected to generate representative transcriptomes of cell type A at time bin 2. Recreation accuracy reflects how closely the model’s output matches this expected outcome.

A high recreation accuracy (approaching 1.0) indicates robust reconstruction of developmental trajectories, while a low accuracy (approaching 0) suggests poor model performance.

Detailed code for generating and evaluating GeneSys-predicted transcriptomes, including calculation of the cell type recreation accuracy, is provided in Jupyter Notebook 4, available in the GitHub repository (see Code Availability section).

### Gene expression programs

Gene expression programs (GEPs) in the root atlas were identified using consensus Non-negative Matrix Factorization tool cNMF (v1.3.1)^26^. The analysis was performed on the raw UMI count matrix using the tool’s default preprocessing steps and parameters. We screened K values ranging from 2 to 50 and selected K = 30, which produced the most stable factorization. cNMF outputs spectra scores for each gene across the 30 GEPs, indicating the strength of association between each gene and each program. In addition, cNMF estimates GEP usage for each cell, quantifying the contribution of each GEP to the cell’s expression profile (Table S1).

Gene ontology (GO) analysis was performed on each GEP using the top 200 genes ranked by spectra score. Enrichment analysis was conducted using gprofiler2 (v0.2.2) and DAVID (https://davidbioinformatics.nih.gov/) with default parameters.

Detail code related to running cNMF and GEP-associated plotting are provided in Jupyter Notebook 8-9, available in the GitHub repository (see Code Availability section).

### Cell type recovery rate

Cell type recovery rate is a metric used to evaluate the influence of specific genes or gene sets on developmental outcomes predicted by GeneSys. It measures the proportion of correctly predicted terminal cell fates based on simulated single-cell transcriptomes generated from an input tensor with masked gene expression, i.e., only the genes of interest retain their expression values, while all other genes are set to zero.

Detail codes related to gene masking, augmentation and cell type recovery rate is provided in Jupyter Notebook 10-11, available in the GitHub repository (see Code Availability section).

### Gold-standard transcription factor list for gene prioritization benchmarking

The gold-standard transcription factor (TF) list used for benchmarking gene prioritization schemes consists of 140 TFs, derived from the intersection of two curated datasets: 1) The MINI-EX list^29^, a published set of 143 TFs experimentally validated to play pivotal roles in root development. 2) A custom StringDB-based list^44^, comprising 209 TFs identified through functional annotations associated with root biology (Table S2).

The StringDB-based TF list was generated by querying the STRING database (https://string-db.org/) for TFs whose gene descriptions or Gene Ontology (GO) terms contained specific root-related keywords. The following keywords were used: root, xylem, phloem, procambium, pericycle, vascular, vasculature, stele, tracheary, sieve, trichoblast, atrichoblast, root hair, epidermis, epidermal tissue, trichome, lateral root cap, cortex, endodermis, ground tissue, columella, quiescent center.

From this list, tissue- and cell type–specific gold-standard subsets were created using more targeted keyword filters: Stele (vascular tissue)-specific: xylem, phloem, procambium. Epidermis-specific: atrichoblast, trichoblast, root hair, trichome, lateral root cap, epidermal tissue. Xylem-specific: xylem, tracheary. Trichoblast-specific: trichoblast, root hair, trichome

### CellOracle and differential expression (DE) analysis for benchmarking

To run the CellOracle pipeline (v0.7.0)^5^, a base gene regulatory network (GRN) was first constructed to represent a comprehensive collection of plausible TF-target interactions. To identify regions of open chromatin, publicly available scATAC-seq data from *Arabidopsis* roots (GSE155304: GSM4698760)^45^ was processed using Cell Ranger ATAC (v1.2.0) to generate a peak-by-cell matrix. Cicero (v1.11.1)^46^ was then used to infer a co-accessibility map of chromatin regions. Transcription start sites (TSS) were annotated using the TAIR10 genome assembly. Peaks with low co-accessibility scores were filtered according to CellOracle’s guidelines (https://morrislab.github.io/CellOracle.documentation/tutorials/base_grn.html).

To enrich the base GRN, we incorporated TF-target interactions from both DAP-seq (DNA affinity purification sequencing)^47^ and a previously established integrative gene regulatory network (iGRN)^48^. The resulting base GRN included a total of 11.7 million interactions, involving 1601 transcription factors and 31019 target genes.

GRNs were then inferred for each of the 36 combinations of cell types and developmental stages present in the wild-type root atlas^12^. The base GRN was restricted to genes that showed dynamic expression along pseudotime in each cell type, along with their associated TFs^49^. Each cell type-specific GRN was constructed using default parameters as recommended in the CellOracle documentation.

To filter GRN edges, the filter_links function was applied, retaining the top 20000 edges with p-values ≤ 0.01 for each subnetwork. Network centrality measures were then computed using CellOracle’s built-in functions. For benchmarking purposes, centrality scores were aggregated across all developmental stages for each gene.

The differentially expressed analysis was performed with Seurat v4 (v4.1.1.9001)^39^ with ROC methods and the following parameters: logfc.threshold = log(2), min.diff.pct = 0.25, max.cells.per.ident = 10000, only.pos = T.

Detailed code for running CellOracle on the wild-type root atlas is available in the GitHub repository associated with our Brassinosteroid GRN publication^25^.

### GeneSys-derived linear interaction matrix (LIMA)

To gain insights into the temporal transcriptomic dynamics learned by GeneSys, we computed Linear Interaction Matrices (LIMAs) that capture gene-gene relationships during state transitions. First, we used the trained GeneSys model to generate representative single-cell transcriptomes, resulting in cell-by-gene (c × g) matrices for specific states (specific cell type and developmental stage/time point). Given two such states, A and B, represented by matrices of gene expression levels across cells, we computed a transformation matrix W (g × g), termed the Linear Interaction MAtrix (LIMA), which describes how gene expression in state A must change to match that in state B.

This relationship is modeled by the linear equation AW = B, where A and B are cell-by-gene expression matrices from the two states. The transformation matrix W is solved as W=A^−1^B. Each entry W_ij_ in the resulting LIMA quantifies the influence of gene j on gene i during the transition: Positive W_ij_ indicates upregulation, negative W_ij_ suggests downregulation, zero indicates no direct interaction, and diagonal entries (W_ii_) represent self-regulation.

The magnitude of each entry reflects the interaction strength. The matrix can be interpreted as a directed, weighted gene interaction network, enabling discovery of key regulators and pathways active during developmental transitions.

### LIMA construction and application for root development

For the *Arabidopsis* root system, we annotated each pseudotime bin with known developmental stages (Figure S5B) and selected five biologically representative state transitions: Stem cell → Proliferation domain (t₀ → t₁). Proliferation domain → Transition domain (t₁ → t₃). Transition domain → Early elongation zone (t₃ → t₅). Early elongation → Late elongation zone (t₅ → t₇). Late elongation zone → Maturation zone (t₇ → t₉).

For each of these five transitions, we derived 10 LIMAs, one for each major cell type, resulting in a total of 50 LIMAs that collectively map the spatiotemporal transcriptional dynamics of the root system. These matrices serve as a foundation for analyzing gene interactions underlying cell fate decisions and developmental progression.

To generate LIMAs, we simulated 2000 cell trajectories, yielding 22k cell transcriptomes across 11 time steps (2000 × 11 = 22k) from an input tensor of shape (2000, 11, 17513), assembled by sampling transcriptomes from the test set. For each transition of interest, we extracted two (200, 17513) matrices representing gene expression in the initial and subsequent states. From these, a (17513 × 17513) LIMA was computed for each cell type and transition. Because the simulation process is stochastic, we repeated it 10 times, generating 10 replicate LIMAs for each transition. To reduce noise and remove outliers, we computed a z-score for each matrix entry across the 10 replicates. Only entries with |z-score| > 3 were retained. For these, the mean value across replicates was used to construct a denoised final LIMA. These denoised LIMAs were then used for gene prioritization benchmarking.

Detailed codes demonstrating how to derive GRNs from GeneSys-generated single-cell transcriptomes and subsequently perform denoising procedures can be found in the Jupyter Notebook number 12 deposited in the GitHub repository (See Codes availability section).

### Network centrality measurements

Network centrality metrics were calculated using the Python package NetworkX v3.1. For each cell state transition, Linear interaction matrices (LIMAs) and CellOracle-inferred gene regulatory networks (GRNs) were analyzed. The following centrality measures were computed for each transcription factor (TF): 1) Betweenness centrality, 2) Eigenvector centrality, 3) Out-degree centrality, 4) In-degree centrality, 5) Total degree (sum of in-degree and out-degree).

To identify candidate tissue- and cell type-specific TFs with elevated activity during state transitions in the root, we applied a gene prioritization strategy based on the combined centrality score, defined as the sum of a TF’s betweenness, out-degree, and in-degree centrality scores. This score was computed across 50 LIMAs, each representing a biologically meaningful state transition.

To evaluate tissue or cell type specificity, we further examined whether the combined centrality score for a particular tissue or cell type contributed more than 50% of the TF’s total centrality score across all transitions. Transcription factors meeting this criterion were considered tissue- or cell type–specific regulators. The resulting candidate TF lists are provided in Table S3.

Detailed code for calculating centrality scores and ranking transcription factors is available in Jupyter Notebooks 12-13 accessible through the GitHub repository (see Code Availability section).

## Codes availability

All source codes, instructions for installation, tutorials, supplementary data and jupyter notebooks documenting the entire analytical processes presented in this study are available at the following GitHub repository: https://github.com/Hsu-Che-Wei/GeneSys

## Acknowledgement

Early reviews of manuscript: Sheng-Yang He, Mingyuan Zhu, Rachel Shahan, Pablo Szekely. Early test of the model: Viktoriia Huryn.

